# Interplay of stability and dynamics in the optimization of a highly proficient *de novo* enzyme

**DOI:** 10.64898/2026.05.29.728685

**Authors:** Sagar Bhattacharya, Gessica M. Adornato, Yuda Chen, Xiao Huang, Wael E. Y. Mouloud, Hyunil Jo, Alexander N. Volkov, Ivan V. Korendovych, Yang Yang, David N. Beratan, Peng Liu, William F. DeGrado

## Abstract

The development of highly active *de novo* enzymes that catalyze new-to-nature reactions is becoming increasingly possible. Here, we identify the features underlying the catalytic efficiency of a highly proficient *de novo* enzyme, from the original computational design to an optimized catalyst obtained through two rounds of directed evolution. Computational, spectroscopic, and biochemical studies reveal successfully designed features, including precise alignment of catalytic residues, transition state stabilization, and environmental tuning. In the most evolved enzyme, binding of a transition state analog also led to widespread increases in backbone conformational stability throughout the protein, except within a helix near the active site entrance, where the introduction of Gly and Pro increased dynamics and catalytic activity. Thus, the entire protein contributes to catalysis in the most optimized enzyme. Also, the initial design considered only the transition state but not substrate binding, leading to a dynamic Michaelis complex prior to optimization. These studies show the multiple features that need to be optimized to achieve high activity in a designed enzyme.

## Introduction

Enzymes are remarkable molecules, capable of catalyzing challenging reactions at high rates and specificities. Traditionally, the mechanisms by which proteins achieve their proficiency^1,2^ have been probed through kinetic^3,4^, spectroscopic and structural studies^5,6^ of native and mutant enzymes^7,8^. While this approach has engendered an understanding of many features that are important for activity, the balance of forces and interactions required to create highly efficient enzymes has remained enigmatic. Moreover, the complex structures, evolutionary histories, and complex biological contexts of natural enzymes make it difficult to discern this balance. Thus, the *de novo* design of enzymes in which the sequence, structure, active sites, and catalytic reactions are not taken from nature represents a critical challenge.

Early attempts at *de novo* design of enzymes focused on redesign of a natural enzyme’s active site to introduce a novel catalytic activity. The activity of these proteins was generally low, and many rounds of mutagenesis were required to achieve enzyme-like catalytic rates^9-14^. More recently, *de novo* enzymes in which the full backbone is also designed from scratch have shown higher initial activity, and it has proven to be easier to optimize their activities to enzyme-like rates^15-19^. Here we examine the structural and dynamic basis for the activity of the *de novo* protein KABLE2.5, which is the most catalytically active *de novo* enzyme lacking metal ions designed to date^17,20^. KABLE2.5 was designed through computational *de novo* design and improved through two rounds of directed evolution. What design principles were responsible for the very high activity of the initial design, and what features were improved through directed evolution? To answer these questions, we studied the mechanism, structure and dynamics of this enzyme family by nuclear magnetic resonance (NMR) and hydrogen-deuterium exchange (HDX), over a time scale spanning nine orders of magnitude from sub-ns to ms.

The Kemp elimination reaction (Fig. 1a) has served for four decades as a target for the design and directed evolution of protein catalysts based on antibodies and computationally redesigned natural enzymes^9-11,16,21-29^, and most recently, fully *de novo* proteins^30^. This reaction involves abstraction of a proton from a C–H bond, similar to enzymes such as triosephosphate isomerase, which uses carboxylate as a general base^31-33^. In our initial design, designated KABLE1, the carboxylate of Asp49 was positioned at the bottom of a deep, dry substrate binding pocket to exclude bulk water, increase contributions of electric fields to catalysis and increase basicity of its carboxylate (Fig. 1a) ^34-37^. The catalytic activity of KABLE1 was improved by a single round of site-saturation mutagenesis from *k*_cat_/*K*_M_ = 460 M^-1^s^-1^ to 230,000 M^-1^s^-1^ in KABLE1.4 (pH 8). An additional round of NMR-guided mutagenesis yielded KABLE2.5 (*k*_cat_/*K*_M_ = 3.2 × 10^6^ M^-1^s^-1^ and *k*_cat_ = 1,400 s^-1^) (Figs. 1b-d) ^17^, whose activity is over an order of magnitude higher than any previous base-catalyzed Kemp eliminases^11,16,29,38^.

**Fig. 1.**
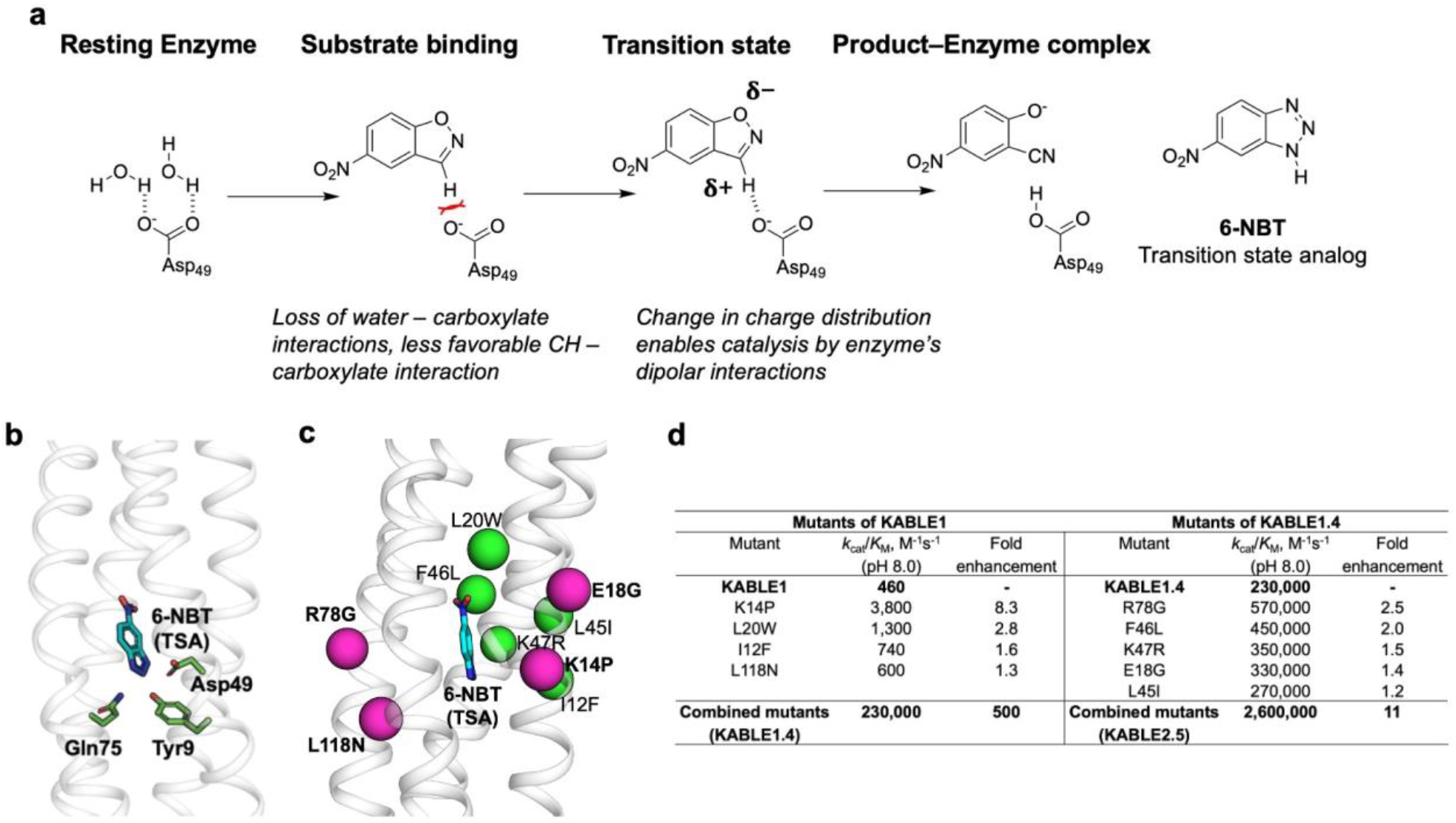
Evolution of *de novo* designed Kemp eliminase KABLE1. **a**, The Kemp elimination reaction, and the transition state analog (TSA) 6-nitrobenzotriazole (6-NBT). Enzymes use part of their substrate-binding energy to create local electrostatic, geometric, and steric frustration, which is relieved as the substrate approaches the transition state. For the Kemp reaction, we targeted the change in partial charge on atoms that occurs as the reaction progresses, for example, converting a relatively apolar aryl C–H of the isoxazole into a more polar group that makes increasingly favorable interaction with Asp49 as the substrate (5-nitrobenzisoxazole, 5-NBI) approaches the transition state. **b**, Key residues in the active site of KABLE1 (gray) are shown as green sticks (TSA shown in cyan). **c**, Beneficial mutations are mapped on the scaffold as magenta (helix-destabilizing mutations) and green colored spheres. **d**, Mutations introduced in first and second rounds of directed evolution are ranked in terms of fold enhancement in *k*_cat_/*K*_M_.

Here, we address the mechanisms by which KABLE1 achieved such a high level of activity, and how it was improved through directed evolution. Two of the substitutions improved hydrogen-bonding, and three other conservative mutants (I12F, L45I, and F46L) optimized packing of the tertiary structure. However, not all the substitutions appeared to be stabilizing; four mutants placed helix-destabilizing residues within alpha-helices (Fig. 1c). In particular, the most active mutant, K14P, placed a helix-breaking Pro near the center of Helix 1 adjacent to the active site, and a second helix-destabilizing mutant, E18G, further increased the catalytic activity of the protein (Fig. 1c).

Here, we show that we got what we designed for in KABLE1. This protein bound the TSA in a hydrophobic pocket precisely as per design. What was not considered but emerged during optimization was tight coupling between the active site and the overall tertiary structure, as well as the structural and dynamic optimization of the Michaelis complex that precedes the transition state complex. As the mutants became catalytically more active, the binding of the TSA became thermodynamically more coupled to the conformational stability of residues throughout the entire protein. One exception was a region surrounding Pro14, which serves as a flexible bridge that becomes increasingly dynamic with increasing activity in KABLE1.4 and KABLE2.5. These findings refine our understanding of fundamental principles of enzymatic catalysis and provide rational targets for design of highly active catalysts.

## Results

### Evaluation of beneficial mutants in two different genetic backgrounds

To understand the effect of substitutions on structure and dynamics it is helpful to first determine whether they act relatively independently or whether they are instead part of a more complex epistatic network. We thus examined the effects of eight productive mutations when introduced into the sequence of KABLE1 versus KABLE1.4 (Fig. 2a and Supplementary Fig. 7, Supplementary Table 8). In each case, individual beneficial mutants introduced into the original KABLE1 sequence had even greater positive effects in the more-evolved KABLE1.4 background sequence. We also have retrospectively noticed this productive combinatorial effect in other *de novo* four-helix bundle enzymes^39,40^. This contrasts with the results of natural proteins, in which many individual beneficial mutations often cannot be combined to create a more active enzyme during evolution^41-43^.

**Fig. 2.**
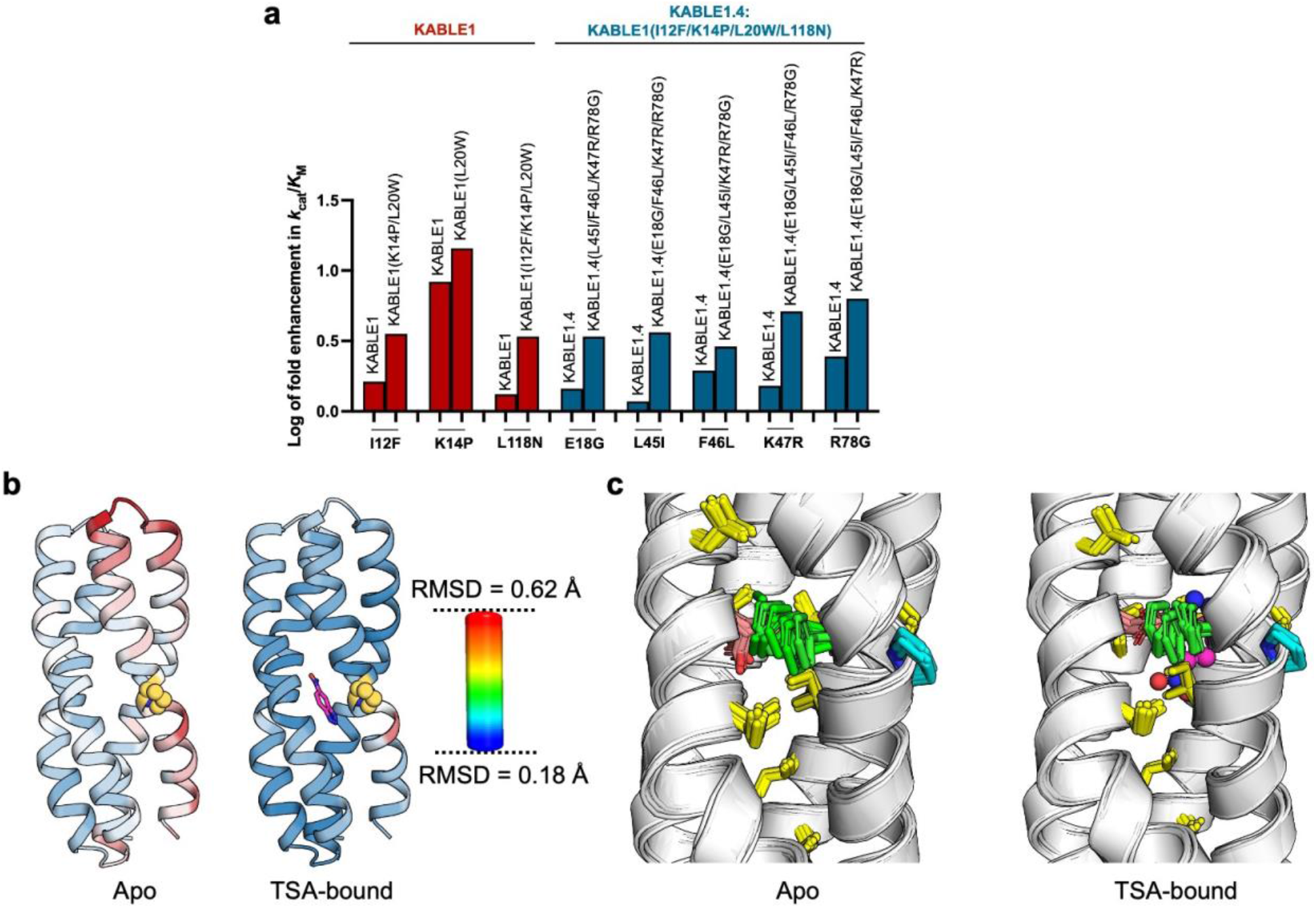
Per-residue RMSD and dynamic fluctuation of I12F in enhancing catalysis. **a**, Log of the fold improvement in catalytic efficiency upon addition of an individual beneficial mutation (shown at the bottom) on: KABLE1 template and combined variants (shown at the top) from first round of directed evolution (red bars); KABLE1.4 template and combined variants (shown at the top) from second round of directed evolution (blue bars). **b**, Per-residue RMSD was mapped onto the apo and TSA-bound structure of KABLE2.5. Per-residue RMSD was computed for each frame using a mean structure generated by averaging coordinates across ten frames of KABLE2.5 NMR structure. The colors range from blue (RMSD = 0.18 Å) to red (RMSD = 0.62 Å). Pro14 is shown with yellow carbon atoms. TSA is shown in magenta carbon atoms. **c**, Dynamic fluctuation of I12F (green sticks) across ten different NMR states for apo (PDB code: 9SMT) and TSA-bound (PDB code: 29SB) structures. Pro14 and Asp49 are shown as cyan and orange sticks, respectively. Hydrophobic packing core residues are shown as yellow sticks.

### NMR measurements show functional tuning of structure, dynamics and stability

The NMR structure of KABLE2.5 in the free and TSA-bound states (Supplementary Table 13) is nearly identical to the original structural model for KABLE1 and crystal structures of proteins of intermediate activity. The structures of KABLE2.5 and its TSA complex were computed from 3600 and 3620 distance restraints, respectively (1423, 1117, and 1060 short, medium, and long-range NOEs, respectively, including 20 intermolecular contacts for the TSA-bound complex, and 237 dihedral angle restraints). The TSA-bound structure is particularly well resolved with an overall backbone root mean square deviation (RMSD) of 0.22 Å, and the free structure is somewhat less well defined, but nevertheless well structured (RMSD = 0.52 Å). Thus, the overall tertiary structure was highly conserved throughout optimization of its activity.

One major exception is the region surrounding Pro14 (Fig. 2b) near the middle of Helix 1, where the chain shows a high RMSD in both the free and TSA-bound states of KABLE2.5. While Pro14 is well ordered, its lack of an amide NH disrupts helical hydrogen bonding. Together, Pro14 and the other substitutions (I12F and E18G; Fig. 2b) appear to induce structural heterogeneity, providing a flexible coupler between the N- and C-terminal halves of Helix 1. We were also interested to find that while the hydrophobic residues at the helix-helix interface were very well-defined, there was a large structural spread in the position of a single side chain, Phe12 (Fig. 2c).

To test the hypothesis that the low helical potential near Pro14 was indeed important for catalysis, we substituted Gly13 and Pro14 with Ala, the most helix-promoting residue of the 20 commonly occurring amino acids^44-46^. Both Gly13 and Pro14 residues are on the surface of the protein and do not directly contact substrate. The P14A mutant had a 6-fold decrease in *k*_cat_/*K*_M_, and the double mutant, G13A/P14A, was 80-fold less active at pH 8. The change in *k*_cat_/*K*_M_ was primarily a result of changes in *k*_cat_ (Supplementary Fig. 6; Supplementary Table 7). These results dramatically illustrate the importance of destabilizing Helix 1 near the active site entrance.

### Dynamics, stability, and thermodynamic coupling between substrate binding and conformational stability of residues distant from the active site

Proteins in solution continuously undergo transient conformational excursions in which subdomains of the protein rapidly equilibrate between the lowest energy “native” structure and a higher energy, more loosely structured state in which the backbone amides are transiently exposed to water. In D_2_O, such transiently exposed residues rapidly exchange with solvent deuterons in a process (called HDX), which can be conveniently monitored by NMR. To provide a measure of stability^47,48^, the rate of HDX (*k*_ex_) at each residue is compared to the intrinsic rate computed for that same residue in an unfolded chain (*k*_int_), yielding position-specific protection factors (PF = *k*_int_/*k*_ex_). The logarithm of PF is a measure of the local conformational stability^47,48^.

While KABLE1, KABLE1.4, and KABLE2.5 in the absence of TSA are hyperstable towards thermal denaturation, their rates of HDX are similar to natural proteins (Fig. 3a). Thus, hyperstability does not imply unusually high rigidity. Protection factors of the helical amides are on the order of 10^4^ at most positions and a small subset of the amides have higher protection factors (> 10^6^). There is no statistically significant difference in the mean of the logarithm of the protection factors for the three proteins. It is also interesting to note that the N-terminal half of Helix 1 and the C-terminal half of Helix 4 have very low protection factors (Fig. 3a). Thus, although these regions, which frame the active site, are well structured in the time-averaged structure of KABLE2.5 (Fig. 2b), they undergo frequent fluctuations in the absence of the TSA. This dynamic behavior might be important for access of substrate and egress of product from the active site, which is largely occluded in the NMR structure of KABLE2.5 and crystal structures of variants of KABLE1.2 (PDB code: 9N0I and 9N0J for apo and TSA-bound protein, respectively).

**Fig. 3.**
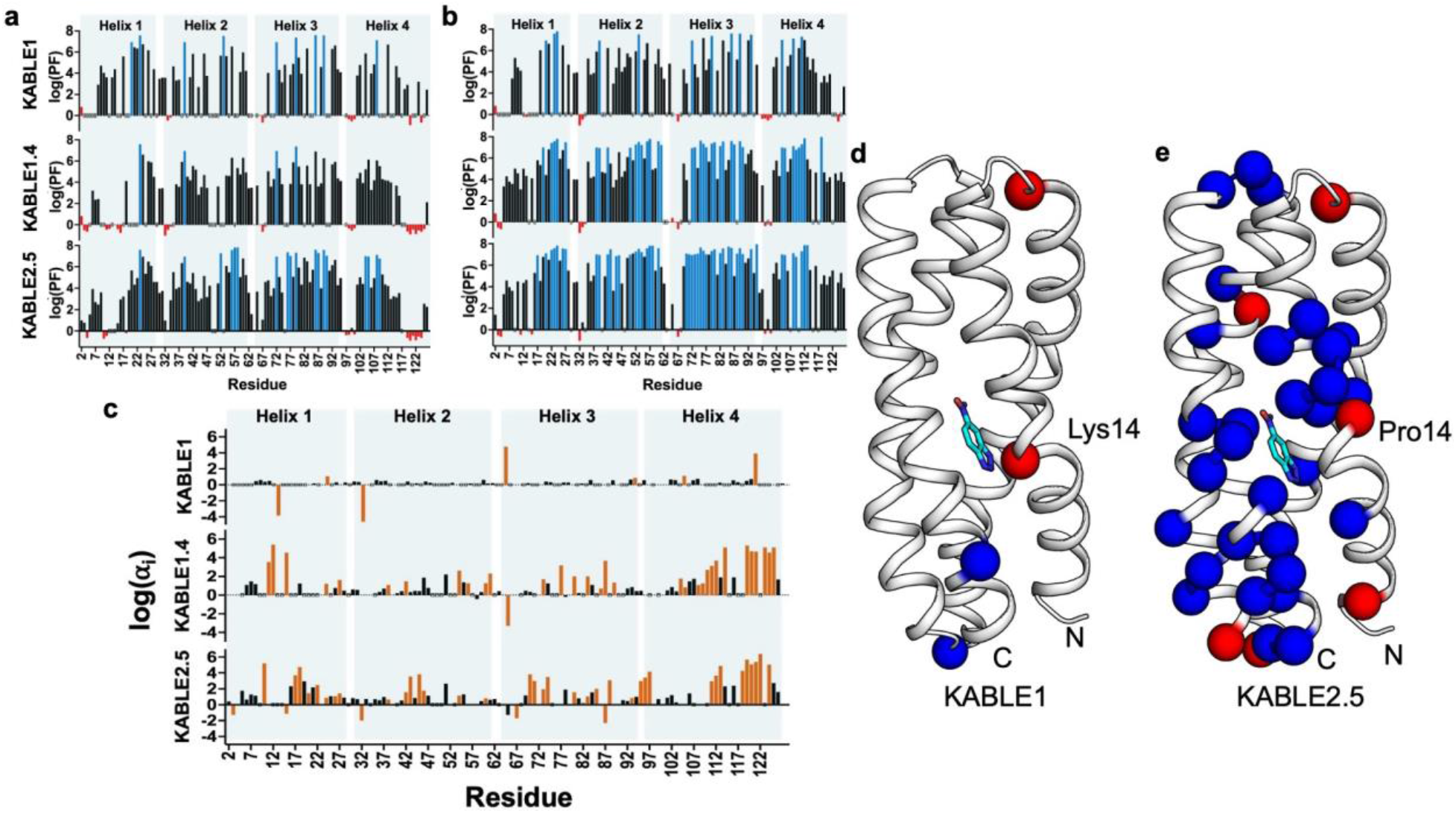
Hydrogen-deuterium exchange of KABLE1, KABLE1.4, and KABLE2.5 backbone amides monitored by NMR (pH 6.9). The PF for each residue was determined by using *k*_int_/*k*_ex_, where *k*_int_ and *k*_ex_ are intrinsic and observed exchange rate, respectively. Values for *k*_int_ were determined by using Sphere^49^, a server program for Hydrogen Exchange Rate Estimation. Log (PF) for **a**, the free proteins and **b**, the TSA-bound proteins obtained by addition of 4 molar equivalents of 6-nitrobenzotriazole. For the resonances that fully exchanged within the measurement deadtime, only a lower limit of *k*_ex_ could be obtained (red). For the signals that showed no decrease in intensity after a week, only an upper limit of *k*_ex_ could be derived (blue). Open floating bar indicates residues where *k*_ex_ could not be obtained. **c**, The log of the fold-increase (log(α_i_)) in protection of KABLE proteins is determined by: PF(bound)_i_/PF(apo)_i_. The mean log(PF) values of the observable residues for 6-NBT-bound KABLE1, KABLE1.4, and KABLE2.5 are 4.501, 5.074, and 5.427, respectively. **d**, log(α_i_) value for each residue for KABLE1 and KABLE2.5 (shown in gray) are mapped on the NMR structure of 6-NBT-bound (shown in cyan) KABLE2.5 (PDB code: 29SB). Blue and red spheres indicate amides that showed a strong increase in protection (log (α_i_) > 2) and strong decrease in protection (log (α_i_) < −1), respectively.

The binding of a ligand to a protein increases its protection factors at sites near the bound ligand. More distant residues will also experience an increase in protection if they are involved in an interaction network that thermodynamically couples them to the binding site. For residues distant from the binding site, the value of log (PF_bound_/PF_apo_)_i_ (called α_i_, where the subscript i references the *ith* residue) is a quantitative measure of the coupling between TSA binding at the active site and the structural stability of residues distant from the binding site^50-52^. Through thermodynamic coupling these same sites must also contribute to the interaction with the ligand.

The value of α_i_ increases at relatively few positions in KABLE1 upon binding of the TSA (Fig. 3b,c), indicating that there is little coupling to residues distant from the binding site in the design. However, substantial increases in α_i_ are seen at numerous sites as one progresses from KABLE1 to KABLE2.5 (Fig. 3c), reaching values of over one million (< −34.9 kJ/mol) at some sites. The sites with very high α_i_ are broadly distributed across the protein in KABLE1.4 and KABLE2.5, showing a large increase in coupling between the proteins. Large changes in properties such as HDX are indeed the hallmark of highly evolved proteins, in which allosteric networks of distant residues contribute to binding and catalysis^53-55^.

While we see large increases in rigidity at most locations in KABLE2.5, there are a few residues where the rigidity decreases instead. Interestingly, in KABLE2.5 one prominent region of low α_i_ occurs near Pro 14 (Fig. 3d,e), indicating that this region is highly flexible. This finding is in excellent agreement with our NMR structural studies, which showed conformational heterogeneity in this region. Other amides with higher α_i_ are at inter-helical turns or the termini of helices where small shifts in the helices can lead to an increase in solvent accessibility of terminal residues.

### Classical molecular dynamics (MD) simulations of KABLE proteins identify regions of increased and decreased dynamics in the most active variant

Given the evidence for dynamics seen by NMR, we turned to MD simulations to evaluate the dynamics of KABLE1, KABLE1.4, and KABLE2.5 on the sub-ns to µs time scale. While previous MD studies of Kemp eliminases have evaluated dynamics near the active sites of apo and TSA complexes^17,56,57^, it is also important to consider the substrate-complexes; thus, we performed triplicate 1-µs simulations of KABLE1, KABLE1.4 and KABLE2.5 in substrate-free, substrate-bound, and TSA-bound states.

Figures 4a-c illustrate profiles of the root mean square fluctuations (RMSF) of the backbone atoms at each position along the protein sequences for the three KABLE variants computed for the TSA-bound, substrate-bound, and substrate-free (apo) states. In the TSA-bound state all three protein structures are considerably less dynamic than the corresponding substrate-free and substrate-bound states. The uniform quenching of dynamics of the TSA-bound structure is consistent with the computational design, which targeted the TSA with the intent of stabilizing the transition state during catalysis. Thus, stabilization of TSA was already achieved in the initial design.

**Fig. 4.**
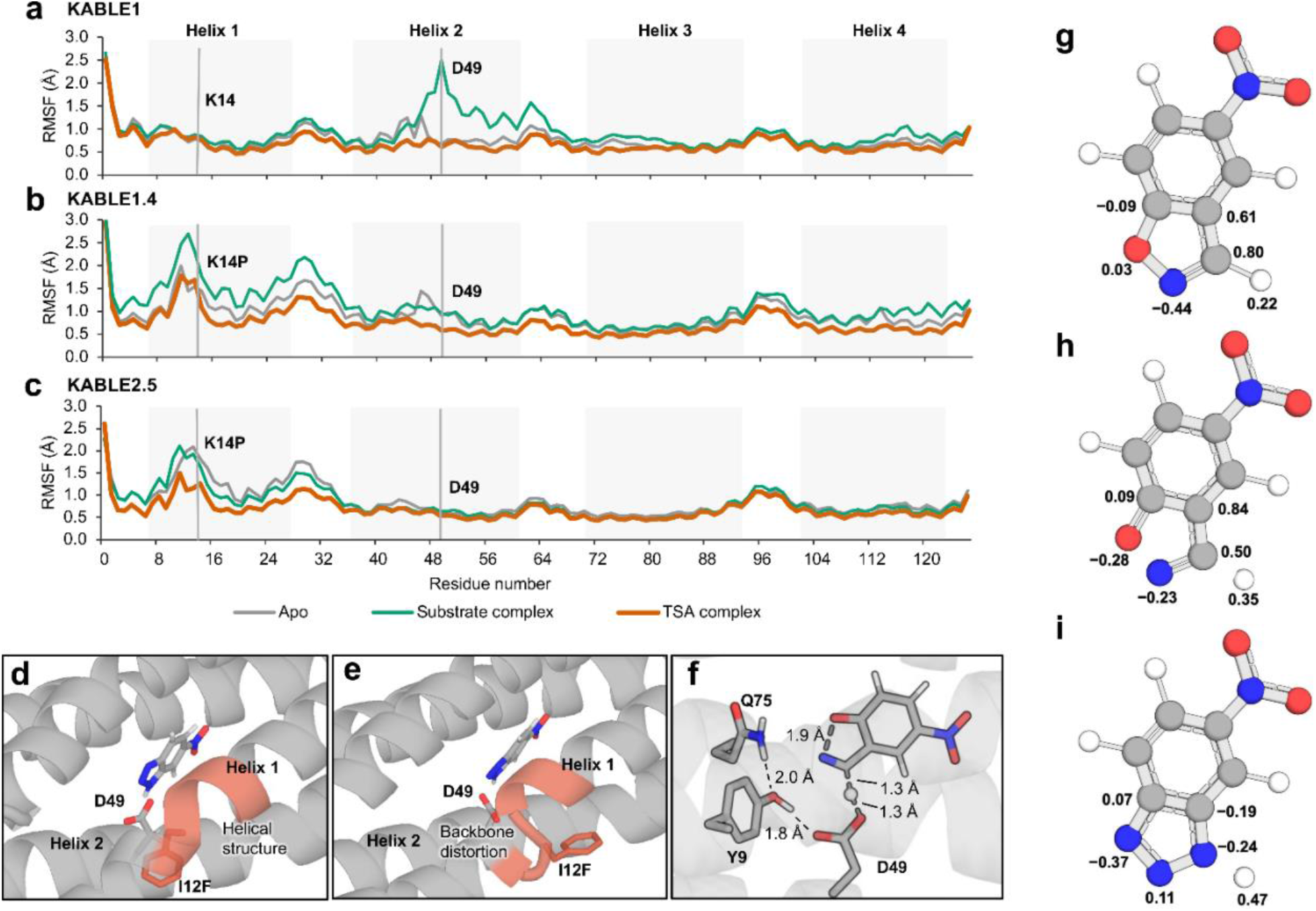
MD simulation and QM/MM analyses of KABLE variants. **(a-c)**, Per-residue RMSFs of backbone heavy atoms of KABLE1, KABLE1.4, and KABLE2.5. The positions of the K14P substitution and Asp49 are indicated by gray lines. Panels **d** and **e** illustrate frames from the simulation of KABLE2.5 complexed with TSA (6-NBT) showing two orientations of Phe12 (I12F). Panel **f** illustrates a QM/MM-computed transition state for the abstraction of a proton from the substrate. QM/MM calculation for charges – **g**, substrate initial state; **h**, substrate transition state; and **i**, TSA.

However, we saw large differences in the position of dynamic regions between the three proteins (Fig. 4a-c). In the apo, substrate-bound, and TSA-bound states, the middle of Helix 1 becomes more dynamic in KABLE1.4 and KABLE2.5 as compared to the parent KABLE1. It is noteworthy that Pro14 is sequentially proximal to two additional helix-breaking Gly residues (^13^GPRILG^18^). Indeed, the helix is locally distorted in this region in KABLE2.5 (Fig. 4d,e).

Just the opposite was seen in Helix 2, which includes both the active site Asp49 as well as several beneficial substitutions (L45I, F46L, and K47R). The least active variant, KABLE1, is uniformly most dynamic in the substrate-bound state, with the largest change near Asp49. Thus, there is a sub-optimal fit with the substrate, which induces more dynamic behavior when bound. In contrast, KABLE2.5 maintains a high degree of stability in Helix 2 in both the apo- and substrate-bound states. KABLE1.4 shows behavior intermediate between KABLE1 and KABLE2.5. Thus, as the catalytic activity increases, there is progressive improvement in the shape complementarity between the protein and the substrate. This finding indicates that one should consider binding of the substrate as well as the transition state when designing enzymes.

We see similar trends when we compare the RMSF over the heavy atoms of the substrate, which vary from 3.5 Å to 2.5 Å to 1.4 Å for KABLE1, KABLE1.4, and KABLE2.5, respectively. Thus, KABLE2.5 is more efficient at aligning the substrate in a pre-attack complex to enter the transition state with minimal change in configurational entropy (Fig. 4f). This computational observation is consistent with the experimental finding that the substrate bound to KABLE1 loses more entropy upon forming the transition state than either KABLE1.4 or KABLE2.5 (see below). We also observed changes in positive partial charge of the atoms within the isoxazole ring of the substrate (Fig. 4g-i), moving from the initial state to the substrate transition state, which are well modeled in the TSA.

We also analyzed the MD trajectories to determine how the active site volume and dynamics are affected upon binding of substrate (shown as histograms from the individual snapshots in Fig. 5a). There is a progressive decrease in active site volume surrounding the TSA progressing from KABLE1 to KABLE2.5, reaching a snug fit for KABLE2.5. We see a similar trend for the substrate complexes, with KABLE2.5 most closely approaching the volume seen when bound to the TSA. Similarly, the active site cavity contracts in each variant in the apo state with KABLE2.5 showing the smallest volume (KABLE2.5 < KABLE1.4 < KABLE1). A large peak is also seen that is associated with a non-productive collapse of the KABLE1.4 active site; this peak is reduced 2.5-fold in KABLE2.5 relative to KABLE1.4. Finally, of the three enzymes, KABLE2.5 most frequently pre-organizes the substrate to move into the transition state by holding the catalytic Asp49 oxygen atom and scissile substrate C–H bond at a distance and angle consistent with a C– H···O hydrogen bond and close to the TS geometry (Fig. 5b-d) ^58^.

**Fig. 5.**
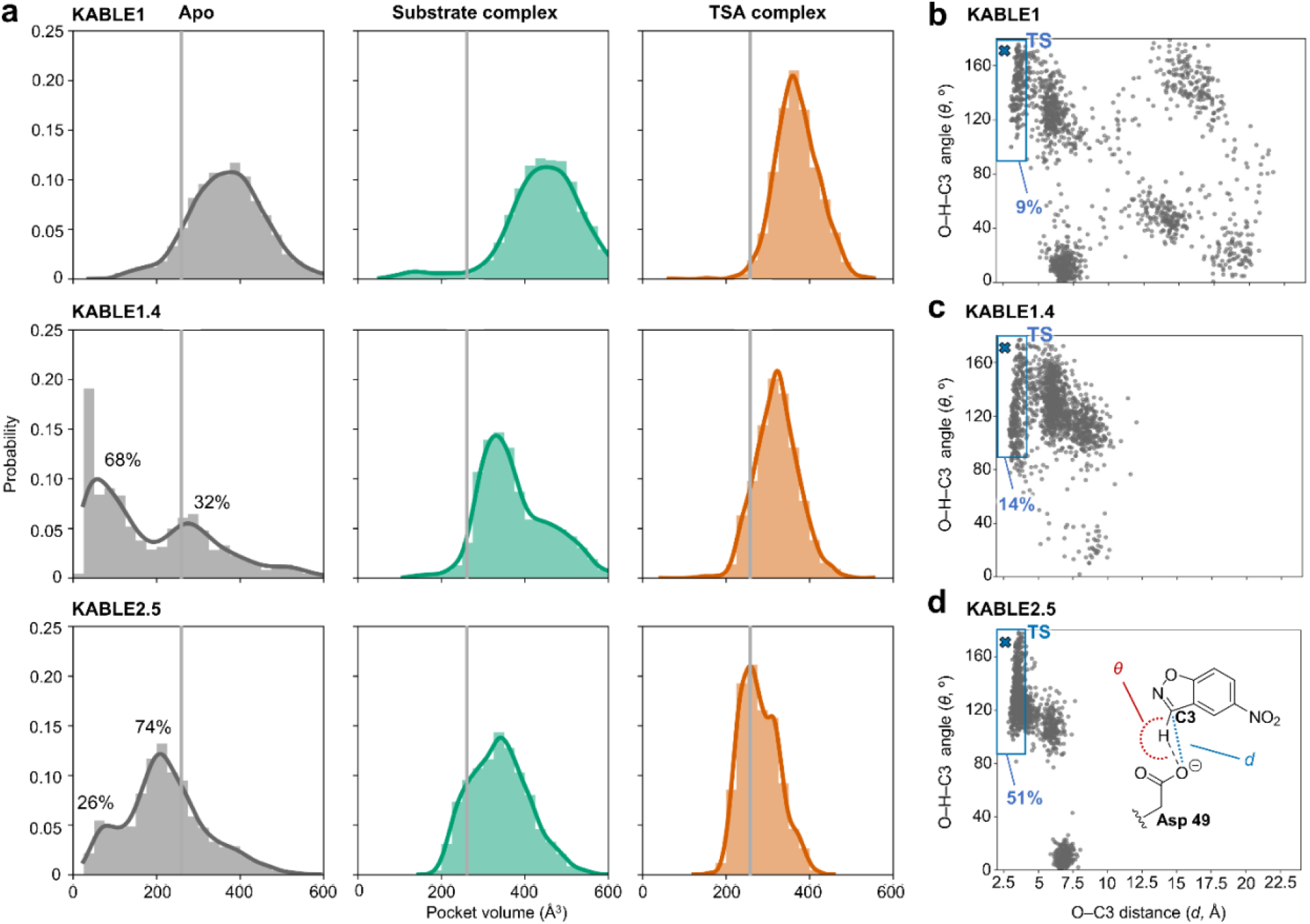
Binding pocket volume histograms for KABLE variants with Kernel Density Estimate fits. **a**, TSA-bound, substrate-bound, and apo proteins. In each case, the gray line indicates the volume observed for the TSA-bound state of the most active variant, KABLE2.5. **(b-d)**, Scatter plots showing the geometry of the substrate with regards to the distance from the D49 carboxylate and the angle of the abstracted C3–H bond for the last 500 ns of simulations with all KABLE variants. The average geometry of the calculated QM/MM transition states is labeled with an “x”. Inset shows a graphical depiction of the plotted distance and angle. A small population of frames with relatively long O–C3 distances are seen in panel b, arising from transient unfolding of the helix near Asp49.

### QM/MM analysis of KABLE2.5 shows the active site residues are well oriented to promote the Kemp reaction

Given that MD simulations revealed progressive substrate preorganization and reshaping of the active site dynamics, we used hybrid QM/MM analysis to assess the reaction path of KABLE2.5 as the substrate is converted through the transition state to the product. The substrate and catalytic residues were treated quantum mechanically and the rest of the protein classically (AMBER ff99SB force field^59^). Five snapshots from an MD trajectory of KABLE2.5 were studied in detail.

The geometry of the transition state for the full protein is similar to that computed in previous fully unrestrained QM analysis of the reacting groups^56,60^. The energetic difference between the substrate and the transition state energy computed for the protein complex was 61.3 kJ/mol, with an associated standard error of 16.4 kJ/mol, which is within error of the experimental value 64.7 ± 0.1 kJ/mol at 25 °C (see below). The derived TS geometry (Fig. 4f) indicates that the protein positions the substrate and Asp49 carboxylate along a low-energy saddle point to promote the carboxylate-catalyzed Kemp elimination reaction. As the transition state is approached, the partial positive charge on the C–H proton increases, producing a greater electrostatic interaction with the active site carboxylate^57,61-63^. We also observe an increase in the partial negative charges on the O atom of the benzisoxazole N–O bond as the substrate approaches the transition state. The carboxamide side chain of Gln75 is well positioned to stabilize this increase in charge.

### Enzyme kinetics of KABLE2.5

Given the large difference in the ordering of the substrate seen in MD simulations, we were motivated to conduct T-dependent kinetics to determine the entropic and enthalpic contributions to catalysis. To interpret these results, we first evaluated the hydrogen-deuterium kinetic isotope effect for the KABLE2.5-catalyzed reaction (by deuterating the CH that is abstracted during elimination). We observed a deuterium isotope effect of 3.8 for *k*_cat_/*K*_M_ for the protein, versus a value of 5–6 for the carboxylate-catalyzed proton abstraction of similar substrates^64,65^. This value suggests that both conformational transitions as well as changes in covalent bonding contribute to the observed rate (Fig. 6a).

**Fig. 6.**
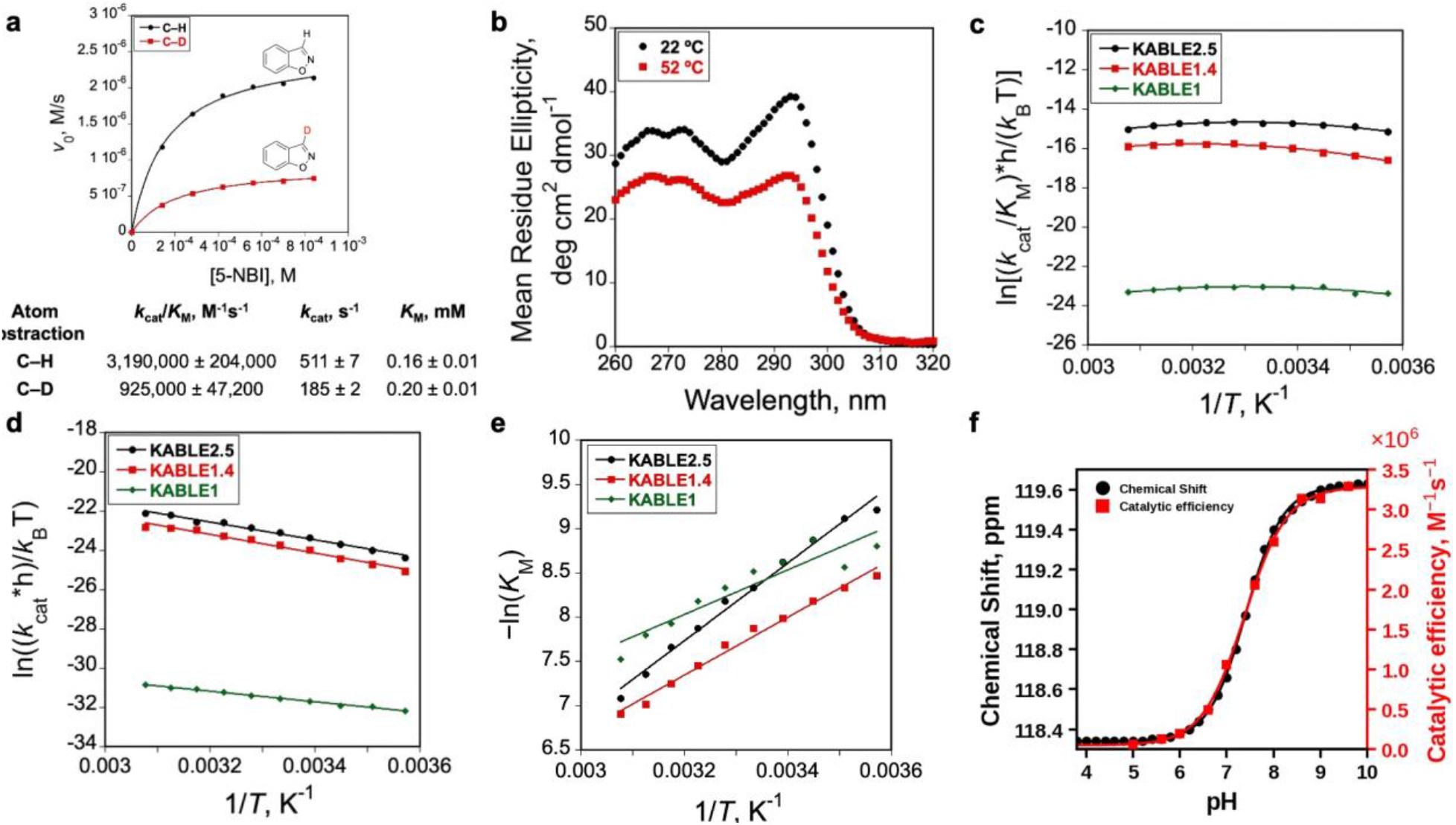
Kinetic isotope effect, pH and temperature dependence, and biophysical characterization of KABLE variants. **a**, Kinetic isotope effects (KIE). Michaelis-Menten plots for KABLE2.5 using protiated (black) and deuterated (red) 5-nitrobenzisoxazole reveal KIEs on *k*_cat_ consistent with turnover being partially limited by product release. Data represent the average of three independent replicates from two different batches of protein. **b**, CD analyses with KABLE2.5 at near-UV region at 22 °C and 52 °C (pH 8.5). Eyring plots for KABLE1, KABLE1.4, and KABLE2.5 at variable temperatures ranging from 7 °C to 52 °C (pH 8.0) – **c**, ln(*k*_cat_/*K*_M_) vs. 1/*T*; **d**, ln((*k*_cat_*h)/*k*_B_T) vs. 1/*T*; and **e**, − ln(*K*_M_) vs. 1/*T*. **f**, pH dependence of chemical shift perturbation (black circle) and catalytic efficiency (red square) of KABLE2.5. Catalytic efficiency at various pH was determined using 5 nM freshly purified KABLE2.5 at 25 °C (pH 8.0) using stopped-flow.

The chemical step (*k*_cat_) of similar C–H deprotonation reactions generally increases with increasing temperature and shows a linear Eyring plot^66^. However, previous temperature-dependent studies of Kemp eliminases have focused on the measurement of *k*_cat_/*K*_M_ rather than evaluating *k*_cat_ and *K*_M_ separately^67,68^, which has resulted in ambiguities in interpretation^69-71^. Thus, to evaluate individual values of *k*_cat_ and *K*_M_, we used stopped-flow kinetics, collecting full substrate concentration/velocity curves at multiple temperatures between 7 °C and 52 °C. This analysis was facilitated by the excellent thermostability of KABLE2.5, as assessed by far- and near-UV circular dichroism (CD) (Fig. 6b, Supplementary Fig. 8), which report on the global stability of secondary and tertiary structure, respectively. For all three KABLE derivatives, we found that *k*_cat_/*K*_M_ has a weak temperature dependence with a slightly domed Arrhenius plot (Fig. 6c), which is a consequence of combining two steps with opposing temperature dependences. By contrast, plots of ln(*k*_cat_) or ln(*K*_M_) versus 1/*T* are both linear (Fig. 6d,e; Supplementary Table 6). The value of *k*_cat_ increases with temperature (Fig. 6d), reaching a rate of 1,680 ± 45 s^-1^ at 52 °C (the highest temperature tested). In contrast, *K*_M_, which is largely a measure of the affinity for the substrate in the Michaelis complex, becomes less favorable with increasing temperature (Fig. 6e).

As the catalytic activity of the KABLE derivatives increases, there is an increasingly unfavorable entropy associated with forming the Michaelis complex; *T*Δ*S* decreases from near 0 for KABLE1 to −15.7 ± 1.4 kJ/mol for KABLE2.5 (evaluated at 298 K, Supplementary Table 6). This finding is consistent with the Michaelis complex having less conformational and configurational entropy, as anticipated in our MD simulations. Also, the opposite trend holds as the substrate moves to the transition state: the magnitude of *T*Δ*S*^‡^ moves from being very unfavorable for KABLE1 (−55.8 ± 0.5 kJ/mol) to much less unfavorable (−19.7 ± 1.6 kJ/mol) for KABLE2.5 (Supplementary Table 6). These data lend credence to our computational observation that the Michaelis complex is structurally better ordered for KABLE2.5 than KABLE1.

Finally, we compared the kinetically determined pH dependence of the enzymatic activity of KABLE2.5 with the corresponding equilibrium value in the absence of substrates. The pH dependence of *k*_cat_/*K*_M_ is well described by a p*K*_a_ value of 7.3 ± 0.1 (Fig. 6f); the corresponding equilibrium p*K*_a_ was 7.4 for substrate-free KABLE2.5, as determined from the NMR chemical shift of Tyr9, an essential active site residue that forms strong hydrogen bonds to Asp49. The good agreement between the kinetically derived and equilibrium p*K*_a_ values supports assignment of the kinetic ionization to Asp49. The active site environment increases the p*K*_a_ of Asp49 by two to three units relative to the intrinsic p*K*_a_ of Asp (approximately 4), making it a better base for proton abstraction.

## Conclusions

An important conclusion of this work is that the entire protein and not just the active site residues contribute to catalysis. We observed the emergence of thermodynamic coupling between binding of the TSA and the conformational stability of regions distant from the binding site during laboratory evolution of KABLE2.5. Because this is a thermodynamically coupled process, it follows that residues distant from the active site contribute to transition state binding, lowering the barrier for the reaction^61,72,73^. Thus, it is noteworthy that distributed allosteric-like and switch-like coupling could be so easily engineered into the KABLE backbone.

Our work also shows the importance of tuning dynamics throughout the protein. Helix 2, which houses the active site residue, became increasingly rigid with increasing catalytic activity. This could be seen experimentally as a decrease in HDX upon binding the TSA; in parallel, MD revealed decreased backbone fluctuations and an increased probability of forming a productive pre-attack geometry in the substrate complex. Consistently, we also observed an increased entropic cost associated with forming the Michaelis complex in the most active variant, which was, however, more than compensated by a decreased entropic penalty associated with moving from the substrate complex to the transition state.

In contrast to the observed increases in rigidity in Helix 2, catalytic activity correlated with increases in dynamics in Helix 1 near the entrance to the active site. The introduction of Pro14 and other helix-disrupting residues resulted in conformational heterogeneity near this residue as seen in the NMR structure; associated increases in local dynamics^74-77^ were seen over a wide range of time scales, from ns fluctuations seen in MD to rarer local unfolding events measured by HDX on ms-to-hour timescales.

These findings provide important lessons for the *de novo* design of enzymes: first, previous attempts to design efficient Kemp eliminases showed that it is much harder to coax an existing natural enzyme to catalyze the reaction by redesigning its active site, than it is to design a *de novo* protein such as KABLE2.5, whose entire structure has been designed to facilitate catalysis. While KABLE2.5 was designed without the use of diffusion algorithms^78,79^, this method is ideal for assuring consistency between form and function. Secondly, we observed the importance of tight coupling between the residues that are close and distant from the active site. Such coupling might be achieved through careful attention to the packing of side chains in concentric rings radiating away from active site residues^54,73,80,81^. Thirdly, by computing MD simulations on the free enzyme, substrate complex, and TSA we learned the importance of considering the full reaction path. While considerable attention has been given to binding transition states and covalent intermediates^11,15,23,60,61^, we found that it is also important to engineer a Michaelis complex to stabilize a productive pre-attack geometry. This should be increasingly possible given the increasing attention to multi-state design^82,83^ and the availability of design tools such as PLACER^15^. Fourthly, we found that, when dynamics were required for function in the KABLE series of proteins, relatively simple substitutions such as helix-breaking residues in Helix 1 were sufficient to deliver high activity. Moreover, Gly and Pro substitutions were found to enhance activity irrespective of whether they were introduced early or late during catalytic optimization, suggesting a rather simple, localized contribution to dynamics and stability. Finally, our findings show that the high thermodynamic stability of the designed proteins to thermal and solvent denaturation does not imply that they are too rigid to enable catalysis.

Together, our findings provide greater insight into Nature’s design of natural enzymes, as well as computational design of novel catalysts. Through the design of proteins we move beyond a passive understanding of enzymes to an active mechanistic understanding that can guide the design of novel catalysts.

## Acknowledgments

We thank members of the DeGrado lab and Liu lab for support.

## Author contributions

S.B., P.L., and W.F.D. formulated the project; S.B. and Y.C. performed cloning, protein expression and purification; S.B. performed directed evolution, kinetic and biophysical characterization. G.M.A. and X.H. conducted MD simulation and QM/MM analyses; W.M. and A.N.V. performed NMR assignment, HDX, and relaxation experiments. S.B., G.M.A., A.N.V., and W.F.D. wrote the manuscript with input from all authors.

## Data, code, and materials availability

All data are available in the main text or in the supplementary information. Chemical shifts (^1^H, ^13^C, and ^15^N) were deposited in the Biological Magnetic Resonance Bank (http://www.bmrb.wisc.edu/) under the accession numbers 53610 (free KABLE1), 53613 (6-NBT-bound KABLE1), 53304 (free KABLE2.5), 53305 (6-NBT-bound KABLE2.5), and 53306 (KABLE2.5 referenced with 7.5 % v/v CD_3_CN). NMR solution structures of free and 6-NBT-bound KABLE2.5 are deposited in the Worldwide Protein Data Bank: wwPDB (https://www.wwpdb.org) under the PDB accession code 9SMT and 29SB, respectively.

## References

1 Knowles, J. R. Enzyme catalysis: not different, just better. Nature 350, 121–124 (1991).

2 Warshel, A. et al. Electrostatic basis for enzyme catalysis. Chem. Rev. 106, 3210–3235 (2006).

3 Fersht, A. Structure and Mechanism in Protein Science (New York: W. H. Freeman, 1999).

4 Kraut, D. A., Carroll, K. S. & Herschlag, D. Challenges in enzyme mechanism and energetics. Annu. Rev. Biochem. 72, 517–571 (2003).

5 Hammes-Schiffer, S. & Benkovic, S. J. Relating protein motion to catalysis. Annu. Rev. Biochem. 75, 519–541 (2006).

6 Fraser, J. S. et al. Hidden alternative structures of proline isomerase essential for catalysis. Nature 462, 669–673 (2009).

7 Bloom, J. D. et al. Thermodynamic prediction of protein neutrality. Proc. Natl. Acad. Sci. U. S. A. 102, 606–611 (2005).

8 Tokuriki, N. & Tawfik, D. S. Protein dynamism and evolvability. Science 324, 203–207 (2009).

9 Rothlisberger, D. et al. Kemp elimination catalysts by computational enzyme design. Nature 453, 190–195 (2008).

10 Khersonsky, O. et al. Bridging the gaps in design methodologies by evolutionary optimization of the stability and proficiency of designed Kemp eliminase KE59. Proc. Natl. Acad. Sci. U. S. A. 109, 10358–10363 (2012).

11 Blomberg, R. et al. Precision is essential for efficient catalysis in an evolved Kemp eliminase. Nature 503, 418–421 (2013).

12 Giger, L. et al. Evolution of a designed retro-aldolase leads to complete active site remodeling. Nat. Chem. Biol. 9, 494–498 (2013).

13 Basler, S. et al. Efficient Lewis acid catalysis of an abiological reaction in a de novo protein scaffold. Nat. Chem. 13, 231–235 (2021).

14 Crawshaw, R. et al. Engineering an efficient and enantioselective enzyme for the Morita–Baylis–Hillman reaction. Nat. Chem. 14, 313–320 (2022).

15 Lauko, A. et al. Computational design of serine hydrolases. Science 388, eadu2454 (2025).

16 Listov, D. et al. Complete computational design of high-efficiency Kemp elimination enzymes. Nature 643, 1421–1427 (2025).

17 Chen, Y. et al. Emergence of specific binding and catalysis from a designed generalist binding protein. Nat. Chem. 18, 1334–1344 (2026).

18 Kim, D. et al. Computational design of metallohydrolases. Nature 649, 246–253 (2026).

19 Braun, M. et al. Computational enzyme design by catalytic motif scaffolding. Nature 649, 237–245 (2026).

20 Davidi, D., Longo, L. M., Jabłońska, J., Milo, R. & Tawfik, D. S. A Bird’s-Eye View of Enzyme Evolution: Chemical, Physicochemical, and Physiological Considerations. Chem. Rev. 118, 8786–8797 (2018).

21 Thorn, S. N., Daniels, R. G., Auditor, M. T. & Hilvert, D. Large rate accelerations in antibody catalysis by strategic use of haptenic charge. Nature 373, 228–230 (1995).

22 Hilvert, D. Critical analysis of antibody catalysis. Annu. Rev. Biochem. 69, 751–793 (2000).

23 Debler, E. W. et al. Structural origins of efficient proton abstraction from carbon by a catalytic antibody. Proc. Natl. Acad. Sci. U. S. A. 102, 4984–4989 (2005).

24 Khersonsky, O. et al. Optimization of the in-silico-designed kemp eliminase KE70 by computational design and directed evolution. J. Mol. Biol. 407, 391–412 (2011).

25 Korendovych, I. V. et al. Design of a switchable eliminase. Proc. Natl. Acad. Sci. U. S. A. 108, 6823–6827 (2011).

26 Privett, H. K. et al. Iterative approach to computational enzyme design. Proc. Natl. Acad. Sci. U. S. A. 109, 3790–3795 (2012).

27 Moroz, Y. S. et al. New Tricks for Old Proteins: Single Mutations in a Nonenzymatic Protein Give Rise to Various Enzymatic Activities. J. Am. Chem. Soc. 137, 14905–14911 (2015).

28 Bhattacharya, S. et al. NMR-guided directed evolution. Nature 610, 389–393 (2022).

29 Gutierrez-Rus, L. I. et al. Enzyme Enhancement Through Computational Stability Design Targeting NMR-Determined Catalytic Hotspots. J. Am. Chem. Soc. 147, 14978–14996 (2025).

30 Beck, J. et al. Customizing the structure of minimal TIM barrels to craft efficient de novo enzymes. Nat. Chem. Biol. (2026).

31 Richard, J. P. A Paradigm for Enzyme-Catalyzed Proton Transfer at Carbon: Triosephosphate Isomerase. Biochemistry 51, 2652–2661 (2012).

32 Clifton, B. E. et al. Evolution of cyclohexadienyl dehydratase from an ancestral solute-binding protein. Nat. Chem. Biol. 14, 542–547 (2018).

33 Deng, H., Dyer, R. B. & Callender, R. Active-Site Glu165 Activation in Triosephosphate Isomerase and Its Deprotonation Kinetics. J. Phys. Chem. B 123, 4230–4241 (2019).

34 Fried, S. D., Bagchi, S. & Boxer, S. G. Extreme electric fields power catalysis in the active site of ketosteroid isomerase. Science 346, 1510–1514 (2014).

35 Schneider, S. H. & Boxer, S. G. Vibrational Stark Effects of Carbonyl Probes Applied to Reinterpret IR and Raman Data for Enzyme Inhibitors in Terms of Electric Fields at the Active Site. J. Phys. Chem. B 120, 9672–9684 (2016).

36 Wu, Y. & Boxer, S. G. A Critical Test of the Electrostatic Contribution to Catalysis with Noncanonical Amino Acids in Ketosteroid Isomerase. J. Am. Chem. Soc. 138, 11890–11895 (2016).

37 Fried, S. D. & Boxer, S. G. Electric Fields and Enzyme Catalysis. Annu. Rev. Biochem. 86, 387–415 (2017).

38 Patsch, D. et al. Enriching productive mutational paths accelerates enzyme evolution. Nat. Chem. Biol. 20, 1662–1669 (2024).

39 Hou, K. et al. De novo design of porphyrin-containing proteins as efficient and stereoselective catalysts. Science 388, 665–670 (2025).

40 Huang, W. et al. De Novo Design, Directed Evolution and Computational Study of Heme-Binding Helical Bundle Protein Catalysts for Biocatalytic Enantioselective Ge–H Insertion. J. Am. Chem. Soc. 147, 40869–40878 (2025).

41 Miton, C. M. & Tokuriki, N. How mutational epistasis impairs predictability in protein evolution and design. Protein Sci. 25, 1260–1272 (2016).

42 Buda, K., Miton, C. M. & Tokuriki, N. Pervasive epistasis exposes intramolecular networks in adaptive enzyme evolution. Nat. Commun. 14, 8508 (2023).

43 Muir, D. F. et al. Evolutionary-scale enzymology enables exploration of a rugged catalytic landscape. Science 388, eadu1058 (2025).

44 O’Neil, K. T. & DeGrado, W. F. A Thermodynamic Scale for the Helix-Forming Tendencies of the Commonly Occurring Amino Acids. Science 250, 646–651 (1990).

45 Brive, L., Dolphin, G. T. & Baltzer, L. Structure and Function of an Aromatic Ensemble That Restricts the Dynamics of the Hydrophobic Core of a Designed Helix-Loop-Helix Dimer. J. Am. Chem. Soc. 119, 8598–8607 (1997).

46 Broo, K. S., Brive, L., Sott, R. S. & Baltzer, L. Site-selective control of the reactivity of surface-exposed histidine residues in designed four-helix-bundle catalysts. Fold. Des. 3, 303–312 (1998).

47 Tsuboyama, K. et al. Mega-scale experimental analysis of protein folding stability in biology and design. Nature 620, 434–444 (2023).

48 Ferrari, Á. J. R. et al. Large-scale discovery, analysis and design of protein energy landscapes. Nature (2026).

49 Zhang, Y.-Z., Roder, H. & Paterson, Y. Rapid amide proton exchange rates in peptides and proteins measured by solvent quenching and two-dimensional NMR. Protein Sci. 4, 804–814 (1995).

50 Englander, S. W. & Kallenbach, N. R. Hydrogen exchange and structural dynamics of proteins and nucleic acids. Q. Rev. Biophys. 16, 521–655 (1983).

51 Englander, S. W., Sosnick, T. R., Englander, J. J. & Mayne, L. Mechanisms and uses of hydrogen exchange. Curr. Opin. Struct. Biol. 6, 18–23 (1996).

52 Englander, S. W., Mayne, L., Kan, Z. Y. & Hu, W. Protein Folding-How and Why: By Hydrogen Exchange, Fragment Separation, and Mass Spectrometry. Annu. Rev. Biophys. 45, 135–152 (2016).

53 Gunasekaran, K., Ma, B. & Nussinov, R. Is allostery an intrinsic property of all dynamic proteins? Proteins 57, 433–443 (2004).

54 Hilser, V. J., Wrabl, J. O. & Motlagh, H. N. Structural and energetic basis of allostery. Annu. Rev. Biophys. 41, 585–609 (2012).

55 Nussinov, R. & Tsai, C. J. Allostery in disease and in drug discovery. Cell 153, 293–305 (2013).

56 Alexandrova, A. N., Röthlisberger, D., Baker, D. & Jorgensen, W. L. Catalytic Mechanism and Performance of Computationally Designed Enzymes for Kemp Elimination. J. Am. Chem. Soc. 130, 15907–15915 (2008).

57 Bhowmick, A., Sharma, S. C. & Head-Gordon, T. The Importance of the Scaffold for de Novo Enzymes: A Case Study with Kemp Eliminase. J. Am. Chem. Soc. 139, 5793–5800 (2017).

58 Panigrahi, S. K. & Desiraju, G. R. Strong and weak hydrogen bonds in the protein-ligand interface. Proteins 67, 128–141 (2007).

59 Hornak, V. et al. Comparison of multiple Amber force fields and development of improved protein backbone parameters. Proteins 65, 712–725 (2006).

60 Na, J., Houk, K. N. & Hilvert, D. Transition State of the Base-Promoted Ring-Opening of Isoxazoles. Theoretical Prediction of Catalytic Functionalities and Design of Haptens for Antibody Production. J. Am. Chem. Soc. 118, 6462–6471 (1996).

61 Frushicheva, M. P., Cao, J., Chu, Z. T. & Warshel, A. Exploring challenges in rational enzyme design by simulating the catalysis in artificial kemp eliminase. Proc. Natl. Acad. Sci. U. S. A. 107, 16869–16874 (2010).

62 Frushicheva, M. P. et al. Computer aided enzyme design and catalytic concepts. Curr. Opin. Chem. Biol. 21, 56–62 (2014).

63 Acosta-Silva, C., Bertran, J., Branchadell, V. & Oliva, A. Kemp Elimination Reaction Catalyzed by Electric Fields. ChemPhysChem 21, 295–306 (2020).

64 Hu, Y., Houk, K. N., Kikuchi, K., Hotta, K. & Hilvert, D. Nonspecific medium effects versus specific group positioning in the antibody and albumin catalysis of the base-promoted ring-opening reactions of benzisoxazoles. J. Am. Chem. Soc. 126, 8197–8205 (2004).

65 Kries, H., Bloch, J. S., Bunzel, H. A., Pinkas, D. M. & Hilvert, D. Contribution of Oxyanion Stabilization to Kemp Eliminase Efficiency. ACS Catal. 10, 4460–4464 (2020).

66 Machado, T. F. G., Gloster, T. M. & da Silva, R. G. Linear Eyring Plots Conceal a Change in the Rate-Limiting Step in an Enzyme Reaction. Biochemistry 57, 6757–6761 (2018).

67 Bunzel, H. A. et al. Emergence of a Negative Activation Heat Capacity during Evolution of a Designed Enzyme. J. Am. Chem. Soc. 141, 11745–11748 (2019).

68 Bunzel, H. A. et al. Evolution of dynamical networks enhances catalysis in a designer enzyme. Nat. Chem. 13, 1017–1022 (2021).

69 Åqvist, J. Computer Simulations Reveal an Entirely Entropic Activation Barrier for the Chemical Step in a Designer Enzyme. ACS Catal. 12, 1452–1460 (2022).

70 Lear, A. et al. Comment on: “Computer Simulations Reveal an Entirely Entropic Activation Barrier for the Chemical Step in a Designer Enzyme”. ACS Catal. 13, 10527–10530 (2023).

71 Åqvist, J. Reply to Comment on: “Computer Simulations Reveal an Entirely Entropic Activation Barrier for the Chemical Step in a Designer Enzyme”. ACS Catal. 13, 10007–10009 (2023).

72 Cooper, A. & Dryden, D. T. Allostery without conformational change. A plausible model. Eur. Biophys. J. 11, 103–109 (1984).

73 Goodey, N. M. & Benkovic, S. J. Allosteric regulation and catalysis emerge via a common route. Nat. Chem. Biol. 4, 474–482 (2008).

74 Woolfson, D. N. & Williams, D. H. The influence of proline residues on alpha-helical structure. FEBS Lett. 277, 185–188 (1990).

75 Woolfson, D. N., Mortishire-Smith, R. J. & Williams, D. H. Conserved positioning of proline residues in membrane-spanning helices of ion-channel proteins. Biochem. Biophys. Res. Commun. 175, 733–737 (1991).

76 Senes, A., Engel, D. E. & DeGrado, W. F. Folding of helical membrane proteins: the role of polar, GxxxG-like and proline motifs. Curr. Opin. Struct. Biol. 14, 465–479 (2004).

77 Chakrabarti, P. & Chakrabarti, S. C--H…O hydrogen bond involving proline residues in alpha-helices. J. Mol. Biol. 284, 867–873 (1998).

78 Watson, J. L. et al. De novo design of protein structure and function with RFdiffusion. Nature 620, 1089–1100 (2023).

79 Ingraham, J. B. et al. Illuminating protein space with a programmable generative model. Nature 623, 1070–1078 (2023).

80 Süel, G. M., Lockless, S. W., Wall, M. A. & Ranganathan, R. Evolutionarily conserved networks of residues mediate allosteric communication in proteins. Nat. Struct. Biol. 10, 59–69 (2003).

81 Motlagh, H. N., Wrabl, J. O., Li, J. & Hilser, V. J. The ensemble nature of allostery. Nature 508, 331–339 (2014).

82 Davey, J. A. & Chica, R. A. Multistate approaches in computational protein design. Protein Sci. 21, 1241–1252 (2012).

83 Friedland, G. D. & Kortemme, T. Designing ensembles in conformational and sequence space to characterize and engineer proteins. Curr. Opin. Struct. Biol. 20, 377–384 (2010).

