## Supplementary Information for "Interplay of stability and dynamics in the optimization of a highly proficient *de novo* enzyme"

Sagar Bhattacharya *et al.*

**The PDF file includes:**

Materials and Methods
Supplementary Figs. 1 to 13
Supplementary Tables 1 to 13
References

**Other Supplementary Materials for this manuscript include the following:**

Movies 1
Supplementary Data 1 to 4

### Supplementary Information

#### Materials and Methods

##### Chemicals and reagents

Buffer salts were purchased from Biobasic, Inc. and Santa Cruz Biotechnology, Inc. Buffers were made using MilliQ water (Advantage A10 System, EMD Millipore).

##### Synthesis of [3-$^{2}$H]-5-nitrobenzisoxazole

The deuterated salicylaldehyde was prepared according to the reported procedure[1] with a yield of 88%. Next, deuterated benzisoxazole was synthesized following Kemp's protocol[2]. 770 mg product (91%) was obtained, which was subjected to nitration according to literature protocols[3,4]. Deuterium incorporation (93%) was determined by $^{1}$H NMR spectroscopy, and the spectrum was consistent with the reported data.

##### Protein expression and purification

The plasmids encoding KABLE variants (Supplementary Table 1), cloned into pET29b(+) vector with NdeI and XhoI restriction sites were obtained from Twist Biosciences, transformed into *E. coli* BL21(DE3) (C2527H, New England Biolabs), and plated on LB agar plates with 50 µg mL$^{-1}$ kanamycin (Thermo Scientific). This same kanamycin concentration was used throughout the experiment. Single colonies were inoculated into 15 mL LB medium containing kanamycin and grown at 37 °C, 220 rpm for 5-6 hours. 10 mL of starter culture was then diluted into 1 L of LB with kanamycin and allowed to grow at 37 °C until $OD_{600}$ reached 0.6-0.8. Culture was induced upon the addition of 0.5 mM isopropyl-β-D-1-thiogalactopyranoside (IPTG) (Apex Bioresearch Products) and grown at 30 °C for 20 hours. Cells were harvested by centrifugation (Sorvall Legend RT+, Thermo Scientific) at 4 °C, 4,150 rpm for 15 min., flash frozen in liquid nitrogen, and stored at −80 °C.

For protein purification, cells were resuspended in buffer A (25 mM Tris, pH 8.0, 20 mM imidazole, 300 mM NaCl). Cells were lysed by sonication (Sonic Dismembrator Model 500, Fisher Scientific) for 10 min. (20 s pulse, 20 s rest) at an amplitude 30%. To isolate the soluble fraction, centrifugation was performed at 4 °C, 20,000*g* for 30 min. The lysate was loaded onto a Ni-NTA gravity column (Hispur, Thermo Scientific) pre-equilibrated with buffer A, washed, and eluted using buffer B (25 mM Tris, pH 8.0, 250 mM imidazole, 300 mM NaCl). Protein fractions identified by Bradford assay were combined, concentrated to 3 mL using 10K MWCO spin concentrator (Amicon Ultra, Millipore Sigma), and exchanged into TEV cleavage buffer (50 mM Tris, pH 8.0, 75 mM NaCl) using desalting column (BioRad, 10 DG). TEV protease (at protein-to-protease absorbance ($A_{280}$) ratio of 100:1) along with EDTA (0.5 mM, pH 8.0, Promega) and DTT (1 mM, Research Products International) was added to cleave the N-terminal $His_6$-tag. Protein was incubated at 37 °C for 12-14 hours, exchanged into buffer C (20 mM HEPES, pH 7.0, 100 mM NaCl). Protein fractions were then passed through Ni-NTA column to remove the protease. Final purification was performed on GE Healthcare AKTA FPLC system with Superdex 75 Increase 10/300 column in buffer C. Protein concentrations were determined by measuring the absorbance at 280 nm using extinction coefficient determined by Expasy ProtParam tool (https://web.expasy.org/protparam/).

##### Expression of $^{13}$C and $^{15}$N labeled protein

For the expression of isotopically labeled KABLE variants, the plasmid was transformed into *E. coli* BL21(DE3) cells and plated on LB agar plate containing kanamycin (50 µg/mL). A single colony was inoculated with 2 mL of LB medium with kanamycin of same concentration mentioned above. The culture was grown at 37 °C for 5-6 hours, then transferred to 18 mL of unlabeled M9 minimal medium with kanamycin and grown further at 37 °C for an additional 5-6 hours. The resulting 20 mL starter culture was diluted with 1 L of M9 medium made with $^{15}NH_4Cl$ and [$^{13}C_6$]-glucose (Cambridge Isotope Laboratories) as $^{15}N$ and $^{13}C$ sources, respectively, supplemented with kanamycin, and grown at 37 °C until $OD_{600}$ reached 0.6-0.8. The culture was then induced by 0.5 mM IPTG and grown at 30 °C for 20 hours. The cells were harvested by centrifugation and the isotopically labeled protein was purified as discussed above.

Kinetic characterization

The catalytic efficiency ($k_{cat}/K_M$) of KABLE variants was determined by measuring initial rates at 380 nm on a stopped-flow spectrophotometer. Product (2-hydroxy-5-nitro-benzonitrile) concentration was determined using an extinction coefficient of 15,800 $M^{-1}cm^{-1}$ at 380 nm. Final substrate 5-nitrobenzisoxazole (5-NBI, AK Scientific) concentrations ranging from 140 μM to 840 μM were used. [3-$^2$H]-5-nitrobenzisoxazole was synthesized and characterized to investigate the primary kinetic isotope effect. The primary substrate 5-NBI and the deuterated version were prepared in acetonitrile as 100 mM stock, which was further diluted in buffer to prepare working solutions of various substrate concentrations. Protein solutions (2x) were prepared in 40 mM Tris, pH 8.0, 100 mM NaCl. Substrate solution (2x) in aqueous solution containing 3% acetonitrile was mixed with either an enzyme solution (containing protein in the buffer) or a control solution (containing only buffer) using KinetAsyst Stopped-Flow System (TgK Scientific) at 25 °C. Upon 1:1 mixing, each control shot contained buffer (20 mM Tris, pH 8.0, 100 mM NaCl), and each enzyme shot contained protein solution (500 nM for KABLE1, 5 nM for KABLE1.4, and KABLE2.5) in buffer (20 mM Tris, pH 8.0, 100 mM NaCl). For all the shots, acetonitrile concentration of 1.5% was maintained as co-solvent. The reaction was followed by monitoring the change in absorbance at 380 nm for 10 s in quintuplicate. Slopes of the linear portion for each individual run from all five measurements were averaged with Kinetic Studio 6 software (v 6.0.5), and after subtracting the mean slope of the control solution set from that of the catalyst solution set, initial rates ($v_0$) were determined. The initial rates, under a series of substrate concentrations in a specified buffer, were used to fit the $k_{cat}$ and $K_M$ using the Michaelis-Menten equation in KaleidaGraph (Synergy Software, v 5.0.6).

pH dependence

The pH dependence of the proteins was performed following the same protocol as described above, except different buffers were tested for different pH (pH 5-5.5: 20 mM citrate, 100 mM NaCl; pH 6.0-9.5: 50 mM Bis-Tris propane, 100 mM NaCl) (Supplementary Fig. 1). Product (2-hydroxy-5-nitro-benzonitrile) concentration was determined using an extinction coefficient of 14,220 $M^{-1}cm^{-1}$ for pH 5.0, while 15,800 $M^{-1}cm^{-1}$ was used for conditions with pH 5.5 and above. The dependence of $k_{cat}/K_M$ on pH was fitted using the equation $k_{cat}/K_M = (k_{cat}/K_M)_{EH} \times (1-f_{E-}) + (k_{cat}/K_M)_{E-} \times f_{E-}$, in which $f_{E-}$ is the fraction of the protein in the anionic state and is related to pH and the p$K_a$ of its base by: $f_{E-}=1/(1+10^{pKa-pH})$ (Supplementary Fig. 2). The $(k_{cat}/K_M)_{E-}$ was used as the maximum $k_{cat}/K_M$. Similarly, $k_{cat} = (k_{cat})_{EH} \times (1-f_{E-}) + (k_{cat})_{E-} \times f_{E-}$. The Michaelis-Menten kinetic parameters are summarized in Supplementary Table 2.

Arrhenius analysis

The temperature dependence of the KABLE series of enzymes was determined by performing Michaelis-Menten kinetic analyses following the same protocol described above except the temperature was varied from 7 °C to 52 °C with an interval of 5 °C (Supplementary Figs. 3-5). The kinetic parameters $k_{cat}$, $K_M$, and $k_{cat}/K_M$ were plotted versus reciprocal of temperature ($T^{-1}$). Data points for $k_{cat}$ and $K_M$ were linearly fitted using the classic Eyring equation (Supplementary Tables 3-5). Thermodynamic parameters are summarized in Supplementary Table 6.

$$\ln\left(\frac{k_{\mathrm{cat}}\,h}{k_B T}\right) = -\frac{\Delta H^{\ddagger}}{R}\left(\frac{1}{T}\right) + \frac{\Delta S^{\ddagger}}{R} \quad (1)$$

$$\ln\left(K_M\right) = \frac{\Delta H^{\circ}}{R}\left(\frac{1}{T}\right) - \frac{\Delta S^{\circ}}{R} \quad (2)$$

where: $k_{cat}$: catalytic turnover
h: Planck's constant
$k_B$: Boltzmann constant
$T$: absolute temperature (K)
R: gas constant
$\Delta H^{\ddagger}$: activation enthalpy
$\Delta S^{\ddagger}$: activation entropy

$K_M$: Michaelis constant
$\Delta H^{\circ}$: Standard enthalpy change for binding
$\Delta S^{\circ}$: Standard entropy change for binding

Circular dichroism

Circular dichroism (CD) spectra of the proteins were recorded using a Jasco J-810 CD spectrometer in continuous scanning mode with 1 nm bandwidth, 1 nm data pitch, scan rate of 50 nm $min^{-1}$. The final spectra represent a buffer-subtracted average of six runs. The spectra of the proteins in the far-UV region (190-250 nm) were collected using a quartz cuvette with a 1-mm pathlength. Protein stocks were diluted in 20 mM HEPES (pH 7.0), 100 mM NaCl buffer to 10 µM, and 300 µL sample was used for analyses. For near-UV region (260-320 nm), 150 µM protein in buffer containing 50 mM Bis-Tris propane buffer (pH 8.5), 100 mM NaCl was used in a cuvette with a pathlength of 10 mm. The mean residue ellipticity (MRE) (in deg $cm^2$ $dmol^{-1}$) values were calculated using the equation: MRE = $\theta/(10 \times c \times l \times N)$, where $\theta$ (mdeg) is ellipticity, $l$ (cm) is pathlength of the cuvette, $c$ (M) is the protein concentration, and $N$ is the number of residues. Thermal denaturation studies were performed by collecting temperature-dependent spectra ranging from 20 °C to 95 °C at 5 °C interval for far-UV region, and 22 °C to 92 °C with 10 °C interval for near-UV region using temperature/wavelength scan mode. MRE values at 222 nm (far-UV) and 295 nm (near-UV) were plotted versus temperature with KaleidaGraph (Supplementary Fig. 8).

Molecular dynamics (MD) simulation

Classical molecular dynamics (MD) simulations were performed for KABLE1, KABLE1.4, and KABLE2.5 for the apo protein, protein–6-NBT complex, and protein–5-NBI complex. Starting structures were derived from the AlphaFold3 server for all simulations of KABLE1. For KABLE1.4 and KABLE2.5, crystal structure of the surface-mutated version of apo KABLE1.2 was used for apo simulation. Similarly, 6-NBT–bound, surface-mutated KABLE1.2 was used for simulation of protein–6-NBT and protein–5-NBI complexes. 5-NBI was docked to 6-NBT ligand by alignment. Surface mutations required for crystallization were mutated back to the kinetically characterized sequence by Rosetta FastRelax. The protonation states of all residues were determined using the H+++ server. The protein and ligands were parameterized using Gaussian 16 and the antechamber program of Amber 24. Ligand partial charges were generated by Restrained

Electrostatic Potential (RESP) charge fitting on geometry-optimized structures using the Merz-Singh-Kollman scheme on electrostatic potentials generated at the B3LYP/6-31g(d) level of theory. All other small molecule parameters were generated by Antechamber using the second-generation general Amber force field (gaff2). The Amber ff19SB force field was used to treat the protein. The simulation box was built by placing the protein in a pre-equilibrated cuboid water box with periodic boundary conditions using the Optimal Point Charge (OPC) water model and 8 Å of padding from the protein. Sodium and chloride ions were added to reach a charge-neutral 100 mM NaCl concentration to match the experimental conditions using the SLTCAP server. Simulations were carried out using the pmemd module of the GPU-accelerated Amber 24 software. Simulations began with 2000 steepest-descent minimization steps followed by a maximum of 8000 conjugate gradient minimization steps. The system was then gently heated to 293 K over 50 ps in the NVT ensemble with a Langevin thermostat and a timestep of 1 fs. During minimization and heating, protein backbone and heavy atoms were restrained with harmonic potentials of 10 kcal/mol•$Å^2$. The system was then equilibrated in the NPT ensemble with a pressure of 1 atm maintained by the Berendsen barostat and a timestep of 2 fs. The restraints on the protein backbone heavy atoms were ramped down from 10 kcal/mol•$Å^2$ to 0 kcal/mol•$Å^2$ over 11 equilibration steps of 100 ps each, totaling 1.1 ns. Finally, unrestrained production simulations were run under the same conditions as equilibration. Three independent replicas were run for each simulation setup. The SHAKE algorithm was used to treat hydrogen atoms, Lennard-Jones and electrostatic interactions cut-offs were set to 8 Å, and long-range electrostatics were calculated using the particle-mesh Ewald method. Subsequent analysis was carried out using the CPPTRAJ module.

Prior to conducting these calculations, we experimentally determined the protonation state of Asp49 using NMR. The pH-rate profile of KABLE2.5 showed a single p$K_a$ of 7.3, which we previously provisionally assigned to Asp49. To evaluate the p$K_a$ of Asp49 at equilibrium, we performed an NMR titration. Direct measurement of Asp49 was complicated by exchange broadening, so we instead measured the transition by monitoring the change in the chemical shift of Tyr9. The chemical shift of Y9's $CH_\varepsilon$ group varied by 0.2 ppm with an apparent p$K_a$ = 7.4. These values are within the range expected for a protonated phenol throughout the titration, strongly suggesting that Asp49 is the ionizable group that is deprotonated at low pH, and that Tyr9 forms a stronger H-bond to the anionic form of D49. QM calculations of chemical shifts were in very good agreement with these assignments and confirmed that D49 was bound in the complex with the transition state analog (TSA).

Quantum Mechanics/Molecular Mechanics (QM/MM) reaction pathway calculations

QM/MM calculations were performed using the ONIOM methodology as implemented in *Gaussian* to investigate the reaction pathway of the KABLE2.5 systems. Initial structural models were obtained from classical molecular dynamics (MD) simulations. From each MD trajectory, one representative snapshot was extracted every 1 ns, yielding five independent frames for each system to sample the conformational variability of the active site.

Prior to QM/MM calculations, all solvent molecules and counter ions were removed from the extracted MD frames. The chosen density functional and basis set combination (B3LYP/6–31+g(d, p) notations in Gaussian16[5]) was selected to appropriately capture polarization effects relevant to an aqueous environment, despite the absence of explicit solvent in the QM/MM calculations.

For each frame, the system was partitioned into three distinct reaction states: **initial state (reactant)**, **transition state**, **product state** (Supplementary Fig. 13, Supplementary Table 9). The QM region consisted of the **substrate**, **Tyr9,** and **Asp49**, which together define the chemically active region. The remaining protein scaffold was treated at the MM level using the

AMBER force field. The QM/MM boundary was defined using standard ONIOM link-atom procedures.
All QM/MM calculations employed electrostatic embedding unless otherwise specified. During geometry optimizations, all atoms in the MM region were held fixed, while all atoms in the QM region were fully relaxed.

Initial-state and product-state geometries were optimized using standard energy minimization protocols. Transition-state optimizations were performed using Gaussian16's[5] transition-state optimization algorithm with analytically computed force constants at the initial step:

```
opt=(calcfc,ts,noeigentest,nomicro,maxstep=15,recalcfc=5)
```

Harmonic vibrational frequency calculations were carried out for all optimized structures. Transition states were confirmed by the presence of a single imaginary frequency corresponding to motion along the reaction coordinate, while initial and product states exhibited no imaginary frequencies. Optimized geometries and QM/MM energies were extracted from the converged calculations for subsequent mechanistic analysis (Supplementary Table 10-11).

<u>Chemical shift calculations</u>

NMR chemical shift calculations were performed to probe electronic perturbations of key active-site residues induced by TSA binding. Two distinct systems were considered: (i) the **free protein**, and (ii) the **TSA–bound complex**.

For the free protein calculations, the QM region consisted of **Tyr9** and **Asp49** only. For the TSA–bound calculations, the QM region included **Tyr9**, **Asp49**, and the **TSA**, which experimentally mimics the transition state geometry of the native substrate. In both cases, the QM/MM boundary was defined at the $\alpha$-carbon of the amino acid residues connecting the QM region to the MM-treated protein environment.

For each system, geometry optimizations and vibrational frequency calculations were first performed at the QM/MM level to ensure well-defined stationary points prior to NMR analysis. Nuclear magnetic shielding tensors were then computed using gauge-including atomic orbital (GIAO) method within the ONIOM framework:

```
oniom(mPW1PW91/6-311+G(2d,p):amber)=embedcharge
nmr(giao)
```

Chemical shifts were referenced against tetramethylsilane (TMS). TMS geometries were optimized at the B3LYP/6-31+G(d,p) level, followed by NMR shielding calculations at the mPW1PW91/6-311+G(2d,p) level to ensure consistency with the QM region treatment:

```
opt freq b3lyp/6-31+G(d,p)
nmr(giao) mPW1PW91/6-311+G(2d,p)
```

For each optimized structure, isotopic chemical shifts were extracted for the **HE1, CE1, HE2, and CE2 atoms of Tyr9** (Supplementary Table 12). Differences in calculated chemical shifts between the free-protein and TSA–bound states were analyzed and compared directly with experimental NMR data to assess the impact of TSA binding on the electronic environment of the active site. The consistency between calculated and spectroscopically measured chemical shifts validates our proposed reaction mechanism in QM/MM calculations.

Nuclear Magnetic Resonance (NMR) spectroscopy

All NMR experiments were performed at 298 K on a Bruker Avance III HD 800 MHz spectrometer equipped with a TCI cryoprobe. For resonance assignments and NMR structure determination, the samples contained 1.4-2.8 mM U-[$^{13}$C, $^{15}$N] labeled KABLE1, KABLE1.4, or KABLE2.5, 20 mM HEPES pH 6.9, 100 mM NaCl, 0.02 % $NaN_3$ and 10 % $D_2O$ for the lock. Samples of proteins bound to the transition state analogue were prepared by a single addition of 4 molar equivalents of 100 mM 6-NBT stock in natural-abundance or perdeuterated acetonitrile (Supplementary Figs. 10-12).

Nearly complete, unambiguous assignments of KABLE1, KABLE1.4, and KABLE2.5 $^{1}$H, $^{13}$C and $^{15}$N backbone resonances were obtained from 2D [$^{1}$H,$^{15}$N] HSQC and a standard set of 3D BEST HNCACB, HN(CO)CACB, HNCO, and HN(CA)CO experiments. For the free and 6-NBT bound KABLE 2.5, the assignments were extended to protein sidechains using 2D [$^{1}$H,$^{13}$C] HSQC and constant-time 2D [$^{1}$H,$^{13}$C] HSQC for the aromatic region; 3D HBHA(CO)NH and (H)CCH-TOCSY experiments; 2D (HB)CB(CGCD)HD and (HB)CB(CGCDCE)HE spectra for the aromatic resonances; and 3D $^{15}$N-edited NOESY-HSQC and $^{13}$C-edited NOESY-HSQC for aliphatics and aromatics. All NMR data were processed in TopSpin 3.6 (Bruker) or NMRPipe[6] and analyzed in CCPN[7]. Assigned $^{1}$H, $^{13}$C and $^{15}$N chemical shifts were deposited in the Biological Magnetic Resonance Bank (http://www.bmrb.wisc.edu/) under the accession numbers 53610 (free KABLE1), 53613 (6-NBT bound KABLE1), 53304 (free KABLE2.5), 53305 (6-NBT bound KABLE2.5), and 53306 (KABLE2.5 referenced with 7.5 % (v/v) $CD_3CN$). NMR assignments of KABLE1.4 were taken from our previous work[8], while those of 6-NBT were obtained from $^{1}$H NMR (800 MHz, 298 K, $CD_3CN$): δ = 8.92 (s, 1H), 8.34 (dd, $^{3}J$ = 9.0 Hz, $^{4}J$ = 1.7 Hz, 1H), 7.99 (d, $^{3}J$ = 9.0 Hz, 1H) and confirmed by the published data[9].

Three-dimensional $^{15}$N-edited NOESY-HSQC and $^{13}$C-edited NOESY-HSQC spectra for aliphatics and aromatics, all acquired with the mixing time of 120 ms, were used for KABLE2.5 structure calculations. For the 6-NBT bound form, the protein NOE dataset was complemented by the intermolecular NOEs obtained from 3D w2-$^{13}$C,$^{15}$N-filtered/edited [$^{1}$H,$^{13}$C] HSQC-NOESY and [$^{1}$H,$^{15}$N] HSQC-NOESY experiments (mixing time 120 ms). The NOE cross-peaks, determined with CCPN Analysis[7], were combined with the dihedral angle restraints, obtained with DANGLE[10], and used as an input for the automated NOE assignment and structure calculations in CYANA v.3[11], followed by the explicit solvent and torsion angle refinement in CNS[12] and Xplor-NIH[13], respectively. The 10 lowest-energy structures were retained and deposited in the PDB under the accession codes 9SMT (free KABLE2.5) and 29SB (6-NBT-bound KABLE2.5). The NMR structure calculation and refinement statistics are presented in Supplementary Table 13. Ramachandran statistics indicate that favored and allowed regions for free KABLE2.5 are 98.3 % and 1.7 %, respectively. The same for 6-NBT–bound KABLE2.5 are 99.0 % and 1.0 %, respectively. No outliers were observed in either of the cases.

NMR pH titrations were performed by an incremental addition of small aliquots (2-5 µL) of 0.5 M NaOH or 0.5 M HCl to the starting samples of 0.7 mM KABLE2.5 or 0.6 mM KABLE2.5 with 4 molar equivalents of 6-NBT in 20 mM HEPES 100 mM NaCl pH 6.9. At each titration point, the pH of the sample was measured with a Spintrode electrode (Hamilton). The pH-dependent spectral changes of Tyr9 Cε/Hε and Asp Cγ resonances were followed, respectively, in a series of constant-time 2D [$^{1}$H,$^{13}$C] aromatic HSQC spectra and carbon-detected 2D CACO experiments with IPAP for virtual decoupling. The pH titration curves were analyzed with a single transition event equation described in the literature[14].

For the hydrogen-deuterium exchange (HDX) experiments, 1 mM U-[$^{15}$N] labeled samples of KABLE1, KABLE1.4 and KABLE2.5 in 20 mM HEPES 100 mM NaCl pH 6.9 were lyophilized and dissolved immediately prior to the measurements in 100 % $D_2O$ (free protein) or $D_2O$ containing 4 molar equivalents of 6-NBT (the bound form). The signal decrease was followed in a series of [$^1H$,$^{15}N$] BEST-HSQC experiments (2 scans, 128 points in the indirect dimension, 0.2 s relaxation delay; total acquisition time of 84 s). The exchange rates were determined by fitting cross peak intensities as a function of time to a single exponential decay curve. Experimental errors were estimated from duplicate measurements performed on independently prepared KABLE2.5 samples, with the exchange rates agreeing to within a factor of two. The HDX rates for fast exchanging KABLE2.5 amides (1–100 $s^{-1}$) were obtained from the CLEANEX experiment[15] acquired with the mixing times of 4, 6, 8, 10, 12, 15, 20, 30, 50, 70, 100, 120, 150, 200, 250, and 300 ms on lyophilized samples dissolved in 100 % $H_2O$ (free protein) or $H_2O$ containing 4 molar equivalents of 6-NBT (the bound form). The CLEANEX signal buildup curves were fitted to a single exponential function as described in the literature[16].

**Supplementary Figures**

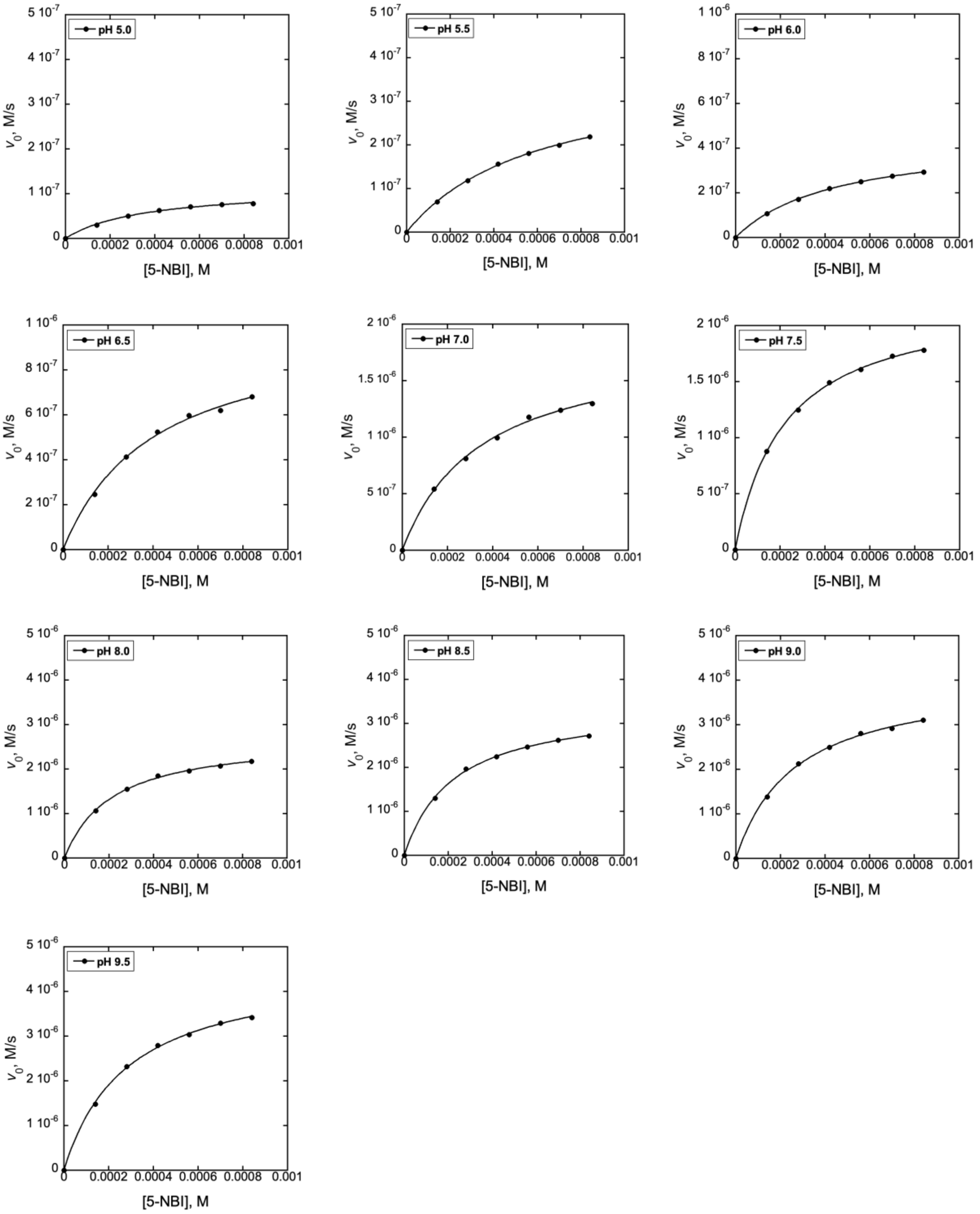


**Supplementary Fig. 1. Michaelis-Menten plots for Kemp elimination catalyzed by KABLE2.5 at different pH.** Final reaction mixtures contained 5 nM enzyme, 140-840 μM substrate 5-NBI, 1.5% acetonitrile in 20 mM Tris, 100 mM NaCl at 25 °C. Kinetic parameters are summarized in

Supplementary Table 2. Data are presented as the mean values and the error bars (most are small compared to the symbol size) represent standard deviations obtained from three independent measurements.

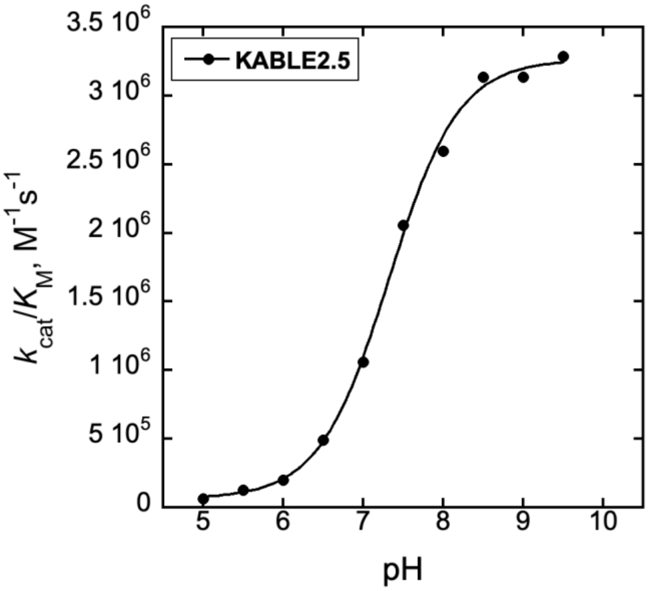


**Supplementary Fig. 2. Dependence of $k_{cat}/K_M$ on pH for KABLE2.5.** Protein concentration – 5 nM. The error bars represent standard deviations in $k_{cat}/K_M$ obtained from three independent measurements, but they are too small to observe compared to the symbol size.

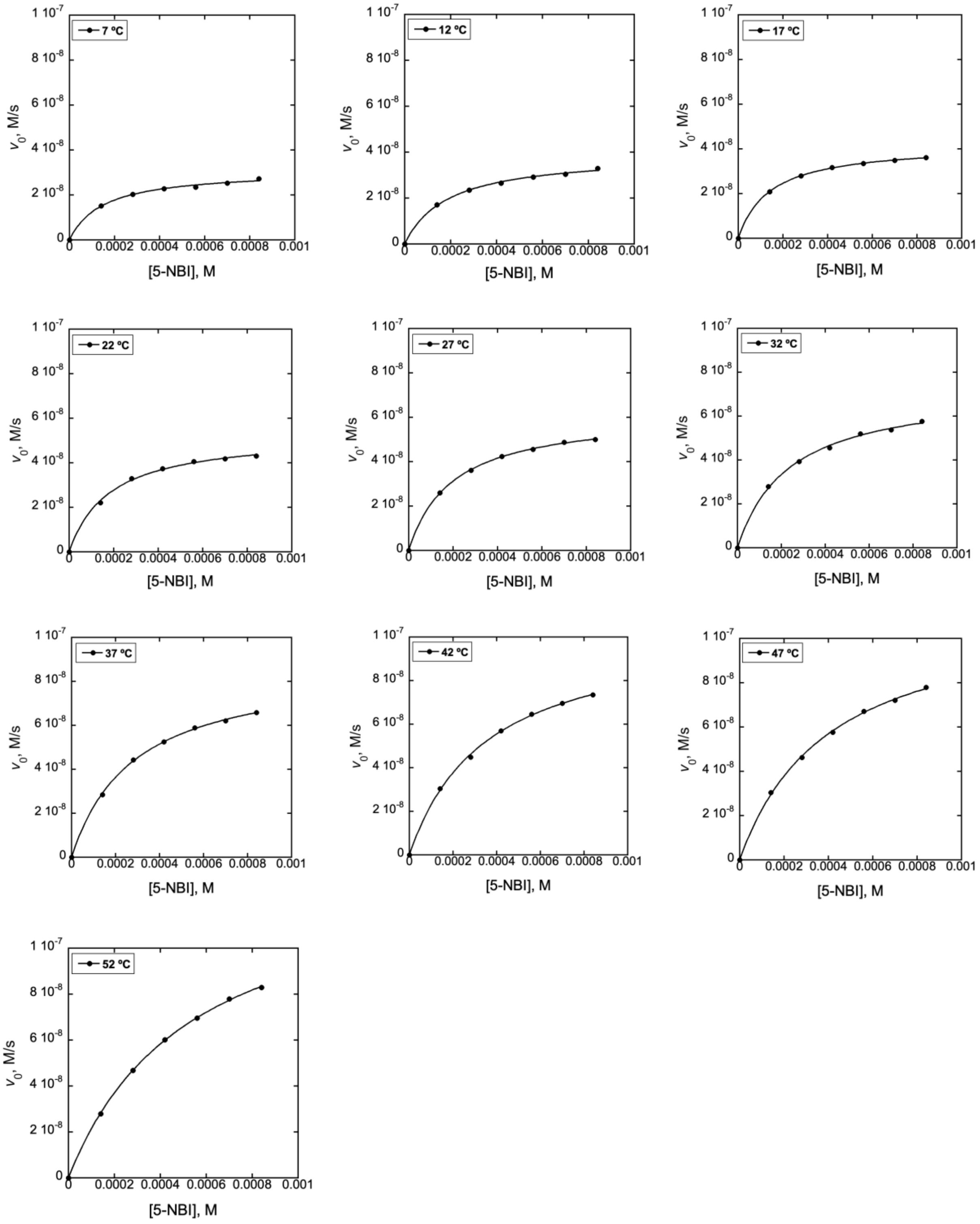


**Supplementary Fig. 3. Michaelis-Menten plots for Kemp elimination catalyzed by KABLE1 at different temperatures.** Final reaction mixtures contained 500 nM enzyme, 140-840 μM substrate 5-NBI, 1.5% acetonitrile in 20 mM Tris (pH 8.0), 100 mM NaCl. Kinetic parameters are summarized in Supplementary Tables 3-5. Data are presented as the mean values and the error bars (most are small compared to the symbol size) represent standard deviations obtained from three independent measurements.

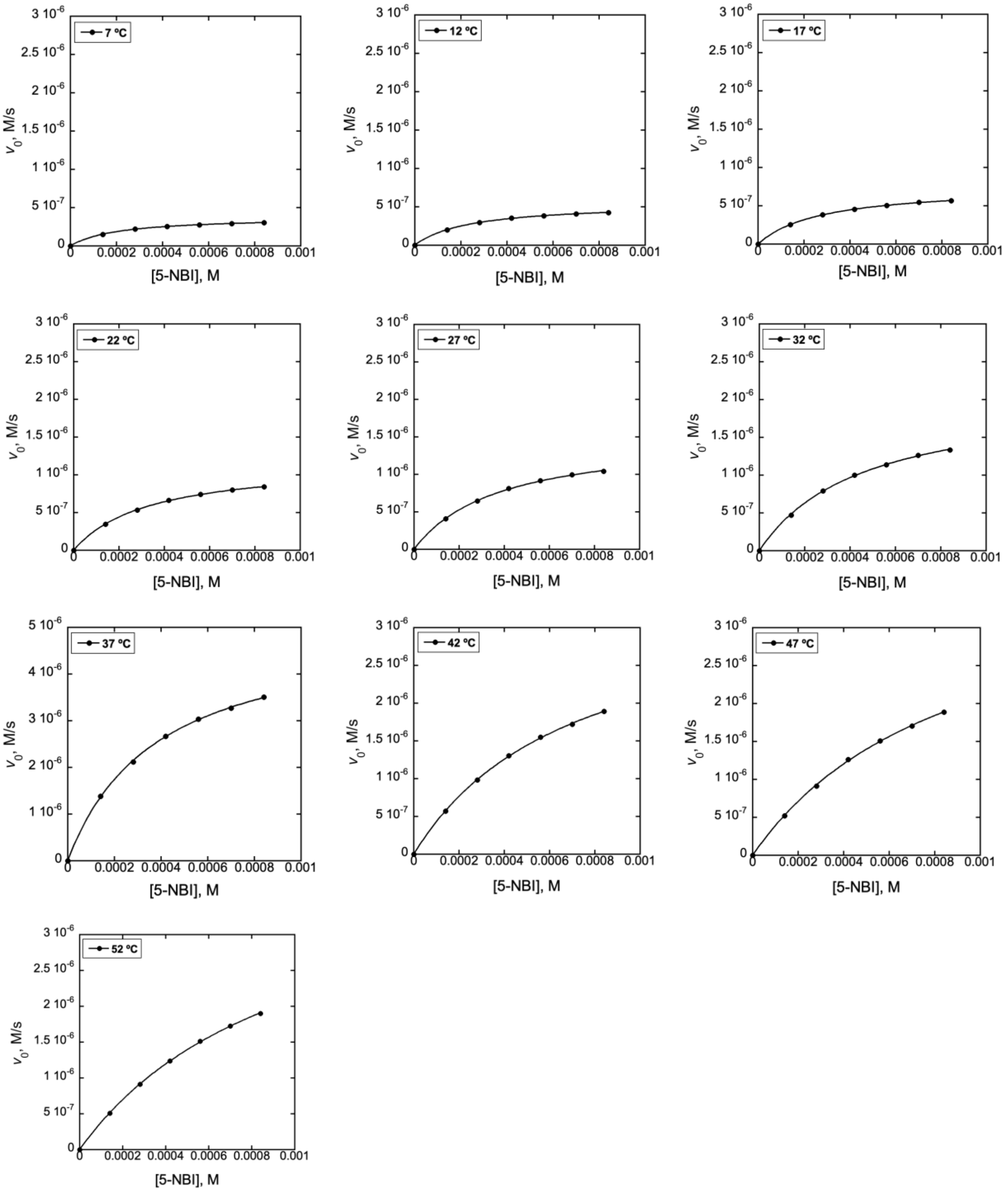


**Supplementary Fig. 4. Michaelis-Menten plots for Kemp elimination catalyzed by KABLE1.4 at different temperatures.** Final reaction mixtures contained 5 nM enzyme, 140-840 $\mu$M substrate 5-NBI, 1.5% acetonitrile in 20 mM Tris (pH 8.0), 100 mM NaCl. Kinetic parameters are summarized in Supplementary Tables 3-5. Data are presented as the mean values and the error bars (most are small compared to the symbol size) represent standard deviations obtained from three independent measurements.

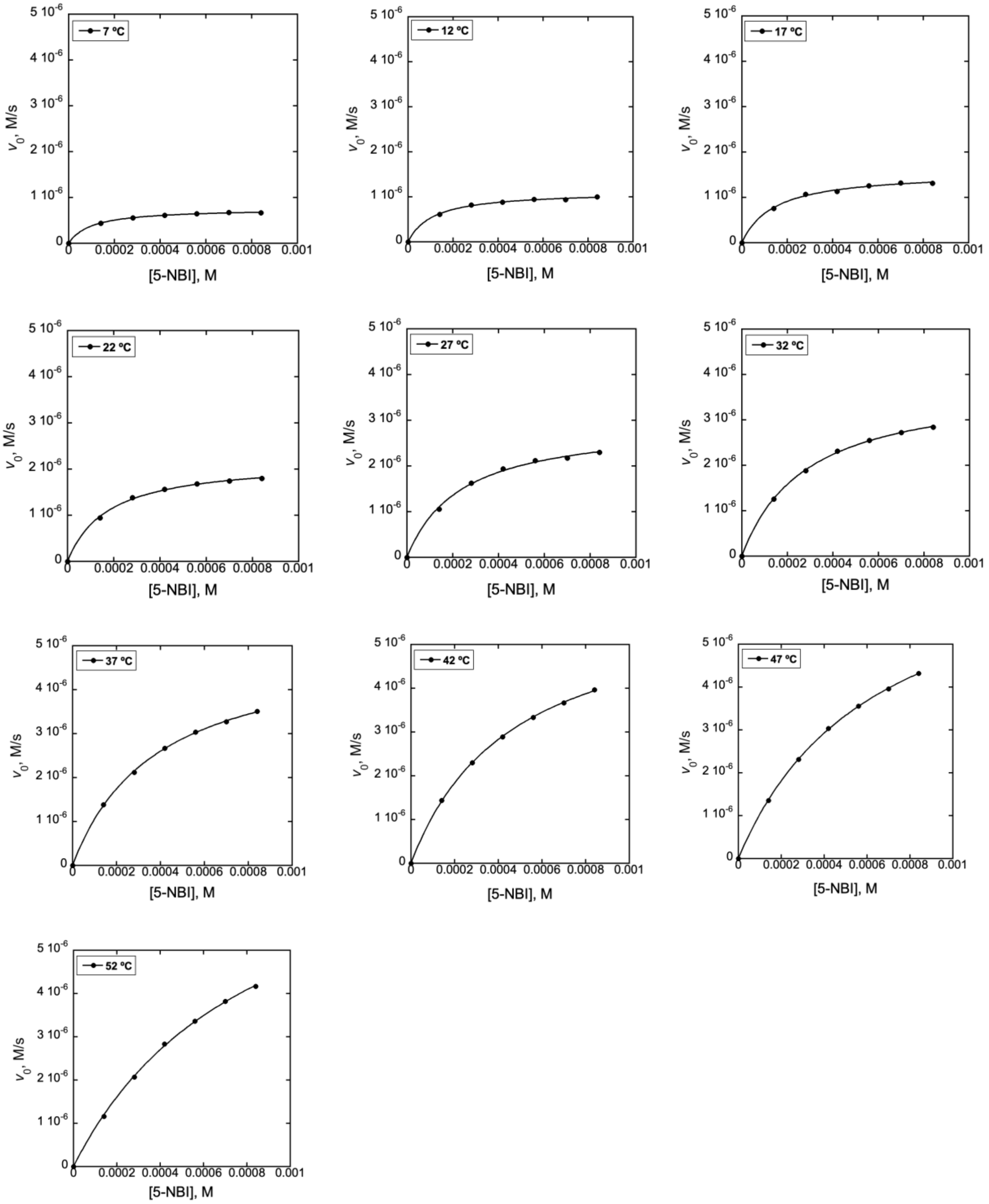


**Supplementary Fig. 5. Michaelis-Menten plots for Kemp elimination catalyzed by KABLE2.5 at different temperatures.** Final reaction mixtures contained 5 nM enzyme, 140-840 μM substrate 5-NBI, 1.5% acetonitrile in 20 mM Tris (pH 8.0), 100 mM NaCl. Kinetic parameters are summarized in Supplementary Tables 3-5. Data are presented as the mean values and the error bars (most are small compared to the symbol size) represent standard deviations obtained from three independent measurements.

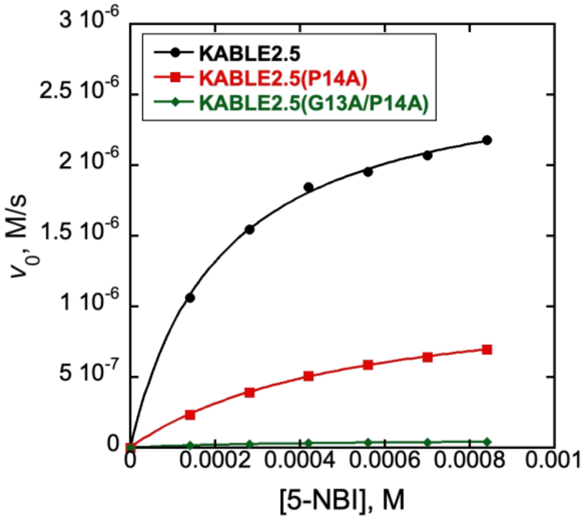


**Supplementary Fig. 6. Michaelis-Menten plots for Kemp elimination catalyzed by P14A and G13A/P14A variants on KABLE2.5 at pH 8.0 (25 °C).** Final reaction mixtures contained 5 nM enzyme, 140-840 μM substrate 5-NBI, 1.5% acetonitrile in 20 mM Tris (pH 8.0), 100 mM NaCl. Kinetic parameters are summarized in Supplementary Table 7. Data are presented as the mean values and the error bars (which are small compared to the symbol size) represent standard deviations obtained from three independent measurements.

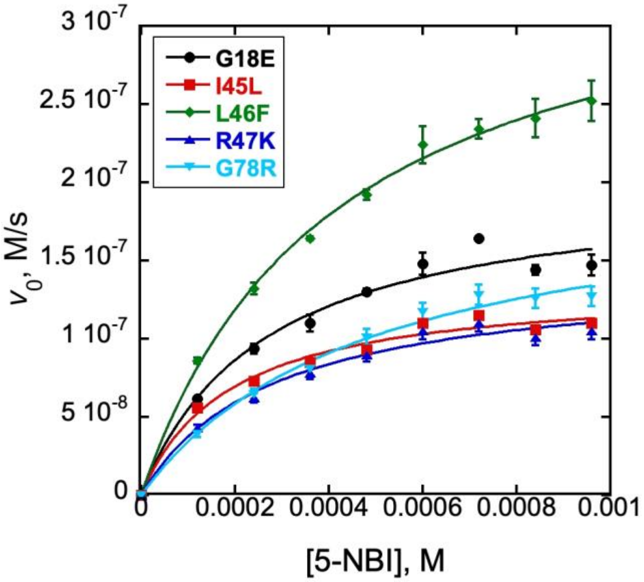


**Supplementary Fig. 7. Michaelis-Menten plots for Kemp elimination catalyzed by individual revertants on KABLE2.5 at pH 8.0 (25 °C).** Final reaction mixtures contained 1 nM enzyme, 120-960 *μ*M substrate 5-NBI, 1.5% acetonitrile in 20 mM Tris (pH 8.0), 100 mM NaCl. Data are presented as the mean values and the error bars represent standard deviations obtained from three independent measurements. Kinetic parameters are summarized in Supplementary Table 8.

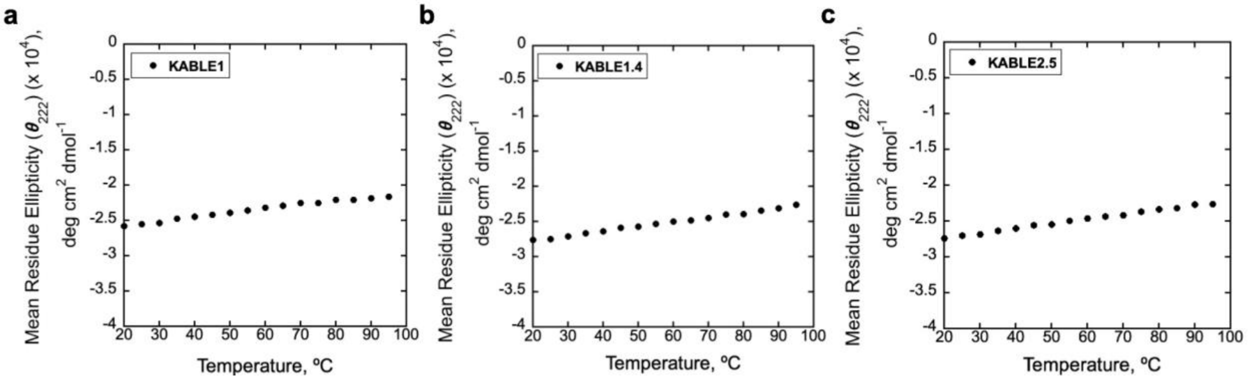


**Supplementary Fig. 8. Thermostability of representative proteins: a, KABLE1; b, KABLE1.4; and c, KABLE2.5 at far-UV region.** CD spectra were collected at 222 nm with 10 μM protein in buffer containing 20 mM HEPES (pH 7.0), 100 mM NaCl.

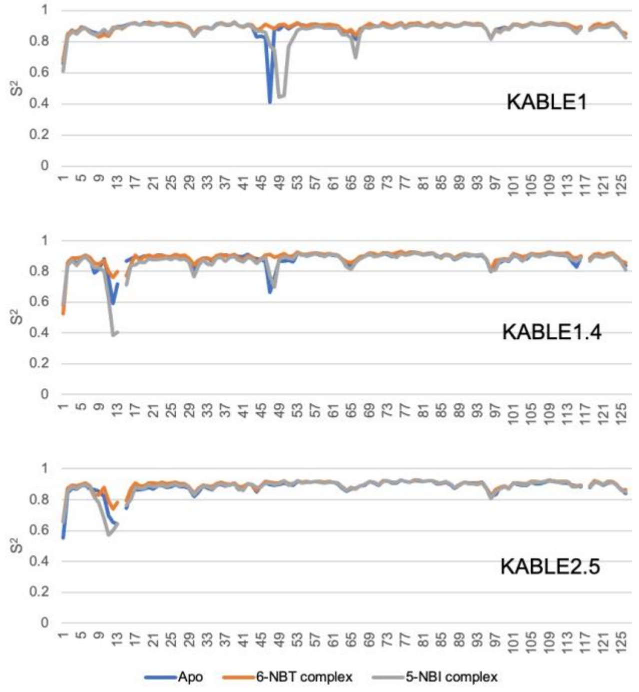


**Supplementary Fig. 9. Order parameters for designed and evolved KABLE variants.**

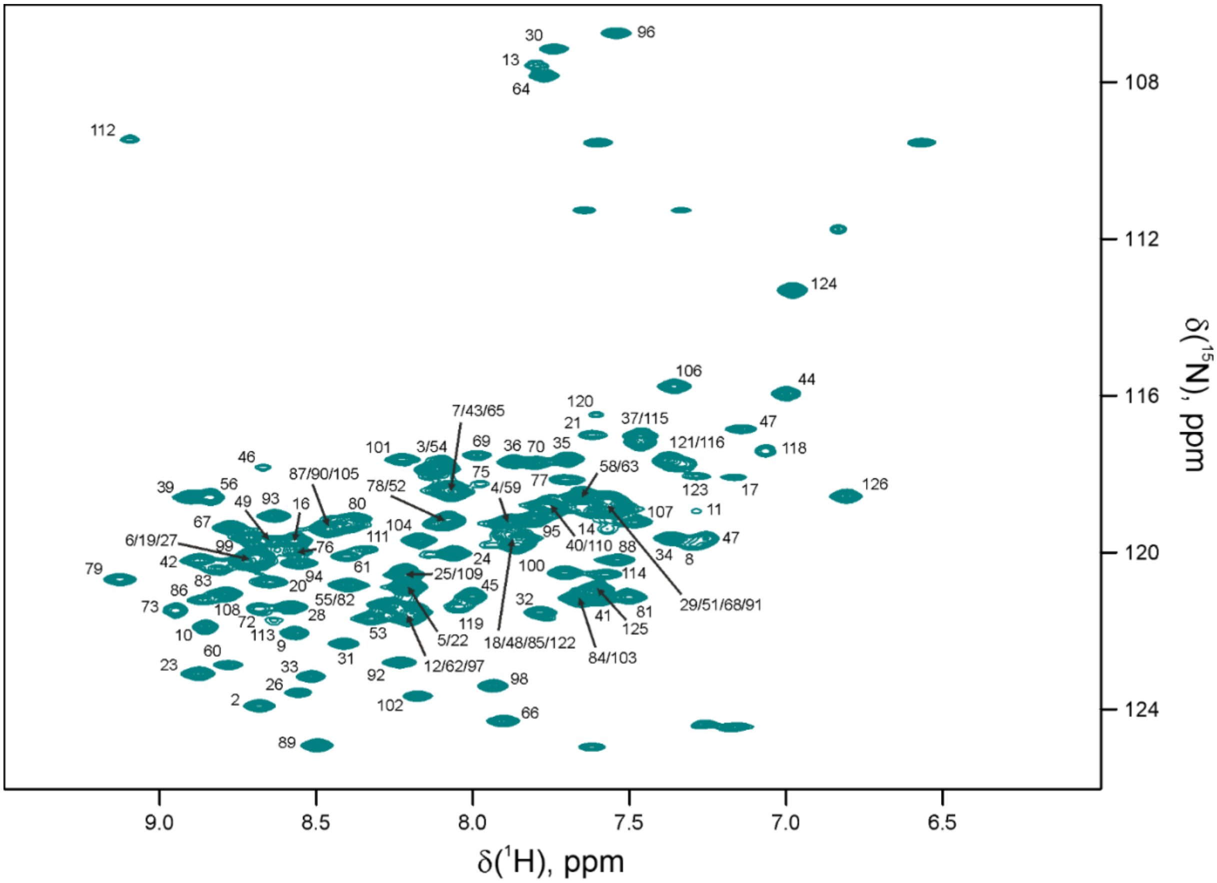


**Supplementary Fig. 10. [$^1$H,$^{15}$N] HSQC spectrum of KABLE1.** Unambiguously assigned backbone amide resonances are labelled with residue numbers. The spectra were recorded in 20 mM HEPES (pH 6.9), 100 mM NaCl at 298 K.

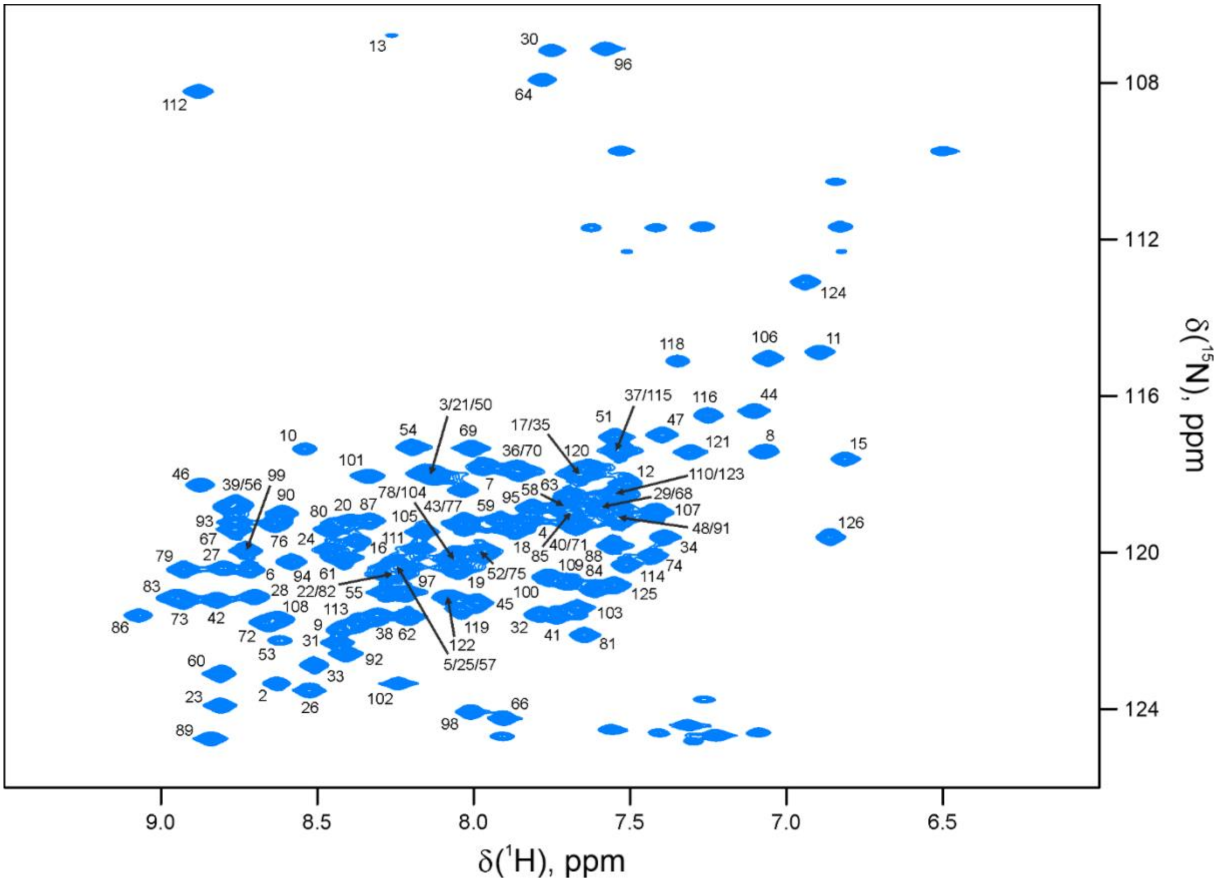


**Supplementary Fig. 11. [$^{1}$H,$^{15}$N] HSQC spectrum of KABLE1.4.** Unambiguously assigned backbone amide resonances are labelled with residue numbers. The spectra were recorded in 20 mM HEPES (pH 6.9), 100 mM NaCl at 298 K.

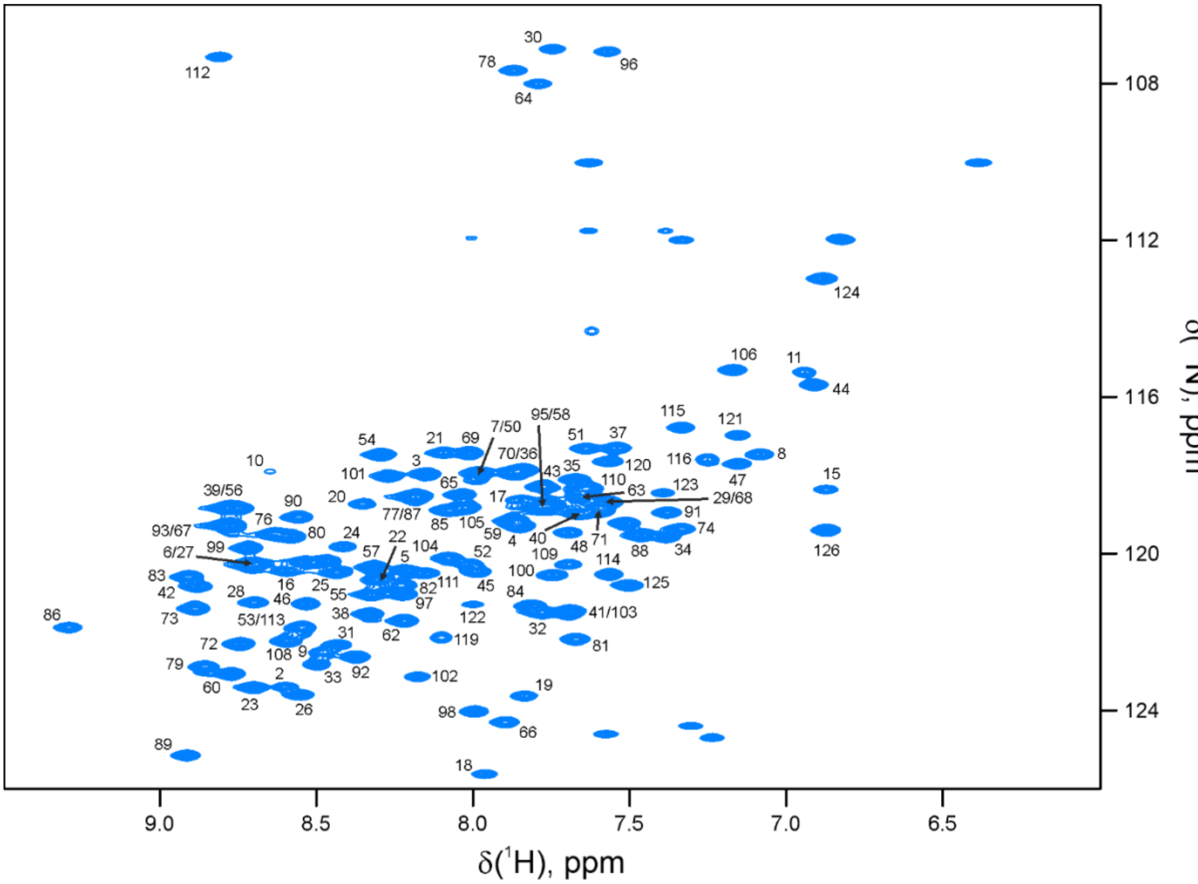


**Supplementary Fig. 12. [$^1$H,$^{15}$N] HSQC spectrum of KABLE2.5.** Unambiguously assigned backbone amide resonances are labelled with residue numbers. The spectra were recorded in 20 mM HEPES (pH 6.9), 100 mM NaCl at 298 K.

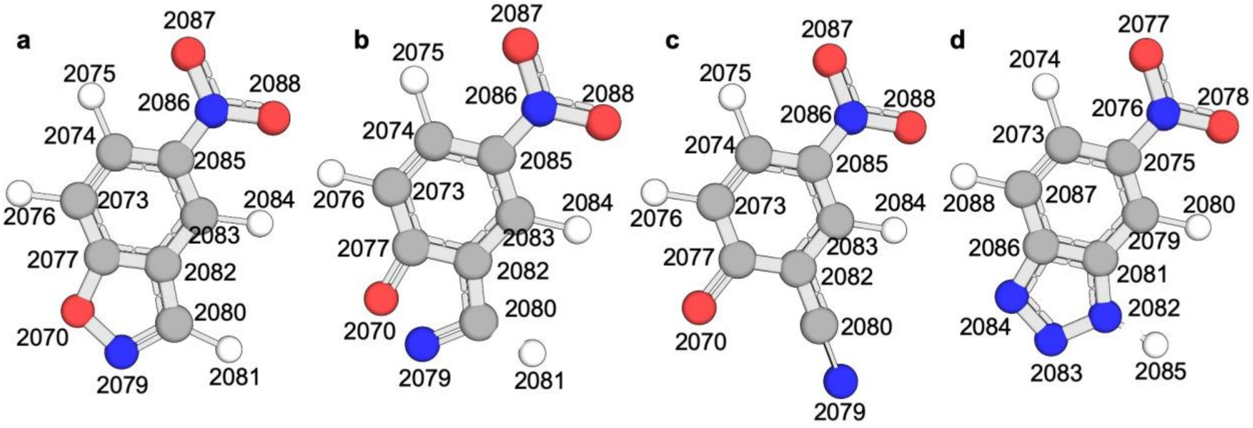


**Supplementary Fig. 13. QM/MM calculation for charges. a,** Substrate initial state; **b,** Substrate transition state; **c,** Substrate product state; and **d,** TSA.

### Supplementary Information

#### Supplementary Tables

**Supplementary Table 1.** DNA and amino acid sequences of the designed and evolved proteins.

| Protein | DNA sequence | Amino acid sequence |
|---|---|---|
| **KABLE1** | ATGCACCACCACCACCACCACGAAAACCTGTACTTCCAGAGCAGCCTCAAAGA<br>AAAGTTTGAGGAGTATGAGAAAATCGGGAAGCGTATTCTTGAACTGTTACAGG<br>AAGCCCGTGACGCTTTCGAGGCTGGTGATCTCGCGCGGGTTGATGAGTTACTT<br>CGGGAATTAAAGGAGCTTTTTAAGAAGGATCTCAAGTTAGCTGAAGAAATGAA<br>AAAGGAAGCGGAAGAAGCTGGGAATAAAGAGGCGGTGGAGCTGCTGGAAGAAC<br>AGCTCGAACGTCTCAAGAAAATCCAGGCGATGTTCGAGGAGGCAGTTGAGGCG<br>TTTCGGGCCGGGGACCGCGAGCGTTTTGGTGAGCTCCTCGAAAAGATCATCGA<br>GGAGGGGAAAGCACTTTTACCTCTTGTGGAGAAGATCAAAGAGGCTATC | MHHHHHHENLYFQSSLKEKFEEYEKIG<br>KRILELLQEARDAFEAGDLARVDELLR<br>ELKELFKKDLKLAEEMKKEAEEAGNKE<br>AVELLEEQLERLKKIQAMFEEAVEAFR<br>AGDRERFGELLEKIIEEGKALLPLVEK<br>IKEAI |
| **KABLE1.4** | ATGCACCACCACCACCACCACGAAAACCTGTACTTCCAGAGCAGCCTCAAAGA<br>AAAGTTTGAGGAGTATGAGAAATTTGGGCCGCGTATTCTTGAACTGTGGCAGG<br>AAGCCCGTGACGCTTTCGAGGCTGGTGATCTCGCGCGGGTTGATGAGTTACTT<br>CGGGAATTAAAGGAGCTTTTTAAGAAGGATCTCAAGTTAGCTGAAGAAATGAA<br>AAAGGAAGCGGAAGAAGCTGGGAATAAAGAGGCGGTGGAGCTGCTGGAAGAAC<br>AGCTCGAACGTCTCAAGAAAATCCAGGCGATGTTCGAGGAGGCAGTTGAGGCG<br>TTTCGGGCCGGGGACCGCGAGCGTTTTGGTGAGCTCCTCGAAAAGATCATCGA<br>GGAGGGGAAAGCACTTTTACCTAACGTGGAGAAGATCAAAGAGGCTATC | MHHHHHHENLYFQSSLKEKFEEYEKFG<br>PRILELWQEARDAFEAGDLARVDELLR<br>ELKELFKKDLKLAEEMKKEAEEAGNKE<br>AVELLEEQLERLKKIQAMFEEAVEAFR<br>AGDRERFGELLEKIIEEGKALLPNVEK<br>IKEAI |
| **KABLE2.5** | ATGCACCACCACCACCACCACGAAAACCTGTACTTCCAGAGCAGCCTCAAAGA<br>AAAGTTTGAGGAGTATGAGAAATTTGGGCCGCGTATTCTTGGTCTGTGGCAGG<br>AAGCCCGTGACGCTTTCGAGGCTGGTGATCTCGCGCGGGTTGATGAGTTACTT<br>CGGGAATTAAAGGAGATTTTGCGTAAGGATCTCAAGTTAGCTGAAGAAATGAA<br>AAAGGAAGCGGAAGAAGCTGGGAATAAAGAGGCGGTGGAGCTGCTGGAAGAAC<br>AGCTCGAAGGTCTCAAGAAAATCCAGGCGATGTTCGAGGAGGCAGTTGAGGCG<br>TTTCGGGCCGGGGACCGCGAGCGTTTTGGTGAGCTCCTCGAAAAGATCATCGA<br>GGAGGGGAAAGCACTTTTACCTAACGTGGAGAAGATCAAAGAGGCTATC | MHHHHHHENLYFQSSLKEKFEEYEKFG<br>PRILGLWQEARDAFEAGDLARVDELLR<br>ELKEILRKDLKLAEEMKKEAEEAGNKE<br>AVELLEEQLEGLKKIQAMFEEAVEAFR<br>AGDRERFGELLEKIIEEGKALLPNVEK<br>IKEAI |
| **KABLE2.5 (P14A)** | ATGCACCACCACCACCACCACGAAAACCTGTACTTCCAGAGCAGCCTCAAAGA<br>AAAGTTTGAGGAGTATGAGAAATTTGGGGCGCGTATTCTTGGTCTGTGGCAGG<br>AAGCCCGTGACGCTTTCGAGGCTGGTGATCTCGCGCGGGTTGATGAGTTACTT<br>CGGGAATTAAAGGAGATTTTGCGTAAGGATCTCAAGTTAGCTGAAGAAATGAA<br>AAAGGAAGCGGAAGAAGCTGGGAATAAAGAGGCGGTGGAGCTGCTGGAAGAAC<br>AGCTCGAAGGTCTCAAGAAAATCCAGGCGATGTTCGAGGAGGCAGTTGAGGCG<br>TTTCGGGCCGGGGACCGCGAGCGTTTTGGTGAGCTCCTCGAAAAGATCATCGA<br>GGAGGGGAAAGCACTTTTACCTAACGTGGAGAAGATCAAAGAGGCTATC | MHHHHHHENLYFQSSLKEKFEEYEKFG<br>ARILGLWQEARDAFEAGDLARVDELLR<br>ELKEILRKDLKLAEEMKKEAEEAGNKE<br>AVELLEEQLEGLKKIQAMFEEAVEAFR<br>AGDRERFGELLEKIIEEGKALLPNVEK<br>IKEAI |
| **KABLE2.5 (G13A/P14A)** | ATGCACCACCACCACCACCACGAAAACCTGTACTTCCAGAGCAGCCTCAAAGA<br>AAAGTTTGAGGAGTATGAGAAATTTGCGGCGCGTATTCTTGGTCTGTGGCAGG<br>AAGCCCGTGACGCTTTCGAGGCTGGTGATCTCGCGCGGGTTGATGAGTTACTT<br>CGGGAATTAAAGGAGATTTTGCGTAAGGATCTCAAGTTAGCTGAAGAAATGAA<br>AAAGGAAGCGGAAGAAGCTGGGAATAAAGAGGCGGTGGAGCTGCTGGAAGAAC<br>AGCTCGAAGGTCTCAAGAAAATCCAGGCGATGTTCGAGGAGGCAGTTGAGGCG<br>TTTCGGGCCGGGGACCGCGAGCGTTTTGGTGAGCTCCTCGAAAAGATCATCGA<br>GGAGGGGAAAGCACTTTTACCTAACGTGGAGAAGATCAAAGAGGCTATC | MHHHHHHENLYFQSSLKEKFEEYEKFA<br>ARILGLWQEARDAFEAGDLARVDELLR<br>ELKEILRKDLKLAEEMKKEAEEAGNKE<br>AVELLEEQLEGLKKIQAMFEEAVEAFR<br>AGDRERFGELLEKIIEEGKALLPNVEK<br>IKEAI |

**Supplementary Table 2.** Kinetic parameters for KABLE2.5 at various pH at 25 °C.

| pH | $k_{cat}$, $s^{-1}$ | $K_M$, mM | $k_{cat}/K_M$, $M^{-1}s^{-1}$ |
|---|---|---|---|
| 5.0 | 23 ± 1 | 0.36 ± 0.03 | 63,900 ± 5,940 |
| 5.5 | 75 ± 2 | 0.60 ± 0.02 | 125,000 ± 5,380 |
| 6.0 | 90 ± 1 | 0.45 ± 0.01 | 200,000 ± 5,000 |
| 6.5 | 202 ± 8 | 0.41 ± 0.04 | 493,000 ± 52,200 |
| 7.0 | 371 ± 11 | 0.35 ± 0.03 | 1,060,000 ± 96,500 |
| 7.5 | 452 ± 4 | 0.22 ± 0.01 | 2,050,000 ± 109,000 |
| 8.0 | 545 ± 8 | 0.21 ± 0.01 | 2,600,000 ± 130,000 |
| 8.5 | 690 ± 4 | 0.22 ± 0.01 | 3,140,000 ± 141,000 |
| 9.0 | 815 ± 13 | 0.26 ± 0.01 | 3,130,000 ± 129,000 |
| 9.5 | 920 ± 15 | 0.28 ± 0.01 | 3,290,000 ± 128,000 |

**Supplementary Table 3.** Catalytic turnover ($k_{cat}$) for designed and evolved KABLE proteins at various temperatures at pH 8.0.

| Temperature, °C | KABLE1 | KABLE1.4 | KABLE2.5 |
|---|---|---|---|
| 7 | 0.062 ± 0.002 | 76 ± 1 | 152 ± 1 |
| 12 | 0.078 ± 0.002 | 110 ± 1 | 224 ± 4 |
| 17 | 0.084 ± 0.001 | 151 ± 1 | 312 ± 8 |
| 22 | 0.106 ± 0.002 | 236 ± 2 | 439 ± 7 |
| 27 | 0.124 ± 0.001 | 305 ± 4 | 593 ± 15 |
| 32 | 0.146 ± 0.003 | 415 ± 7 | 765 ± 8 |
| 37 | 0.176 ± 0.002 | 511 ± 7 | 1,019 ± 12 |
| 42 | 0.210 ± 0.004 | 698 ± 9 | 1,227 ± 14 |
| 47 | 0.230 ± 0.005 | 783 ± 20 | 1,520 ± 14 |
| 52 | 0.274 ± 0.004 | 834 ± 15 | 1,680 ± 45 |

**Supplementary Table 4.** Michaelis constant ($K_M$) for designed and evolved KABLE proteins at various temperatures at pH 8.0.

| Temperature, °C | KABLE1 | KABLE1.4 | KABLE2.5 |
|---|---|---|---|
| 7 | 0.15 ± 0.02 | 0.21 ± 0.01 | 0.10 ± 0.01 |
| 12 | 0.19 ± 0.01 | 0.24 ± 0.01 | 0.11 ± 0.01 |
| 17 | 0.14 ± 0.01 | 0.28 ± 0.01 | 0.14 ± 0.01 |
| 22 | 0.18 ± 0.01 | 0.34 ± 0.01 | 0.18 ± 0.01 |
| 27 | 0.20 ± 0.01 | 0.38 ± 0.01 | 0.24 ± 0.02 |
| 32 | 0.24 ± 0.02 | 0.46 ± 0.02 | 0.28 ± 0.01 |
| 37 | 0.28 ± 0.01 | 0.58 ± 0.02 | 0.38 ± 0.01 |
| 42 | 0.36 ± 0.02 | 0.71 ± 0.02 | 0.47 ± 0.01 |
| 47 | 0.41 ± 0.02 | 0.90 ± 0.04 | 0.64 ± 0.01 |
| 52 | 0.54 ± 0.01 | 1.00 ± 0.03 | 0.84 ± 0.04 |

**Supplementary Table 5.** Catalytic efficiency ($k_{cat}/K_M$) for designed and evolved KABLE proteins at various temperatures at pH 8.0.

| Temperature, °C | KABLE1 | KABLE1.4 | KABLE2.5 |
|---|---|---|---|
| 7 | 413 ± 58 | 362,000 ± 18,900 | 1,520,000 ± 152,000 |
| 12 | 411 ± 24 | 458,000 ± 19,700 | 2,040,000 ± 189,000 |
| 17 | 600 ± 43 | 539,000 ± 59,300 | 2,230,000 ± 129,000 |
| 22 | 589 ± 35 | 694,000 ± 20,800 | 2,440,000 ± 141,000 |
| 27 | 620 ± 32 | 803,000 ± 23,300 | 2,470,000 ± 215,000 |
| 32 | 608 ± 52 | 902,000 ± 38,800 | 2,730,000 ± 101,000 |
| 37 | 629 ± 24 | 881,000 ± 32,600 | 2,680,000 ± 77,800 |
| 42 | 583 ± 34 | 983,000 ± 30,500 | 2,610,000 ± 62,700 |
| 47 | 561 ± 30 | 870,000 ± 43,500 | 2,380,000 ± 42,800 |
| 52 | 507 ± 12 | 834,000 ± 29,200 | 2,000,000 ± 110,000 |

**Supplementary Table 6.** Thermodynamic parameters (from Eyring equation) for designed and evolved KABLE proteins at pH 8.0 (The entropy is evaluated at 298 K).

| Parameter for activation energy | KABLE1 | KABLE1.4 | KABLE2.5 |
|---|---|---|---|
| $\Delta H^{\ddagger}_{\mathrm{cat}}$ (kJ mol$^{-1}$) | 22.47 ± 0.54 | 39.54 ± 1.92 | 37.95 ± 1.59 |
| $T\Delta S^{\ddagger}_{\mathrm{cat}}$ (kJ mol$^{-1}$) | −55.81 ± 0.54 | −19.71 ± 1.92 | −19.71 ± 1.55 |
| **Parameter for binding** | **KABLE1** | **KABLE1.4** | **KABLE2.5** |
| $\Delta H_{\mathrm{bind}}$ (kJ mol$^{-1}$) | −20.96 ± 2.80 | −27.20 ± 1.09 | −36.53 ± 1.42 |
| $T\Delta S_{\mathrm{bind}}$ (kJ mol$^{-1}$) | −0.13 ± 2.76 | −7.74 ± 1.09 | −15.69 ± 1.42 |
| **$\Delta G^{\ddagger}_{\mathrm{overall}}$ (kJ mol$^{-1}$)** | **57.40 ± 4.02** | **39.83 ± 3.14** | **36.82 ± 2.97** |

**Supplementary Table 7.** Michaelis-Menten Kinetic parameters for alanine variants on KABLE2.5 at pH 8.0 (25 °C).

| Enzyme | $k_{cat}$, s$^{-1}$ | $K_M$, mM | $k_{cat}/K_M$, M$^{-1}$s$^{-1}$ |
|---|---|---|---|
| **KABLE2.5** | **545 ± 8** | **0.21 ± 0.01** | **2,600,000 ± 130,000** |
| KABLE2.5 (P14A) | 226 ± 3 | 0.52 ± 0.01 | 435,000 ± 10,000 |
| KABLE2.5 (G13A/P14A) | 12 ± 1 | 0.37 ± 0.06 | 32,400 ± 5,840 |

**Supplementary Table 8.** Michaelis-Menten Kinetic parameters for individual revertants on KABLE2.5 at pH 8.0 (25 °C).

| Enzyme | $k_{cat}$, $s^{-1}$ | $K_M$, mM | $k_{cat}/K_M$, $M^{-1}s^{-1}$ |
|---|---|---|---|
| G18E | 201 ± 16 | 0.26 ± 0.06 | 773,000 ± 186,000 |
| I45L | 135 ± 7 | 0.19 ± 0.03 | 711,000 ± 121,000 |
| L46F | 363 ± 12 | 0.41 ± 0.03 | 885,000 ± 70,800 |
| R47K | 142 ± 9 | 0.28 ± 0.05 | 507,000 ± 96,400 |
| G78R | 202 ± 15 | 0.49 ± 0.08 | 412,000 ± 74,200 |
| **KABLE2.5** | **545 ± 8** | **0.21 ± 0.01** | **2,600,000 ± 130,000** |

**Supplementary Table 9.** Summarized partial charge information for substrate including average partial charge values with standard deviations across different states.

| Atom number\ State | Initial State | Standard deviation | Transition State | Standard deviation | Product State | Standard deviation | TSA | Standard deviation |
|---|---|---|---|---|---|---|---|---|
| 2073,C | 0.2528 | 0.1701 | -0.0118 | 0.1778 | 0.1281 | 0.1847 | 0.1620 | 0.1130 |
| 2074,C | -0.0945 | 0.2473 | 0.0036 | 0.1903 | 0.1864 | 0.2609 | -0.0448 | 0.0762 |
| 2075,H | 0.1545 | 0.0050 | 0.1473 | 0.0133 | 0.1424 | 0.0059 | 0.1542 | 0.0110 |
| 2076,H | 0.1585 | 0.0095 | 0.1387 | 0.0145 | 0.1286 | 0.0152 | 0.1373 | 0.0085 |
| 2077,C | -0.0852 | 0.1746 | 0.0874 | 0.1670 | -0.4525 | 0.1107 | 0.0653 | 0.1533 |
| 2078,O(2084,N for TSA) | 0.0288 | 0.0140 | -0.2833 | 0.0407 | -0.5768 | 0.0111 | -0.3677 | 0.0981 |
| 2079,N | -0.4369 | 0.0315 | -0.2338 | 0.0645 | -0.5669 | 0.0597 | 0.1101 | 0.0523 |
| 2080,C(2082,N for TSA) | 0.7961 | 0.2131 | 0.5053 | 0.1815 | 0.3742 | 0.2080 | -0.2411 | 0.0323 |
| 2081,H | 0.2174 | 0.0064 | 0.3472 | 0.0096 | 0.4130 | 0.0281 | 0.4725 | 0.0084 |
| 2082,C | 0.6139 | 0.2255 | 0.8399 | 0.4848 | 0.8283 | 0.2286 | -0.1915 | 0.2394 |
| 2083,C | -1.0271 | 0.1077 | -1.0459 | 0.1935 | -0.4032 | 0.2341 | -0.1102 | 0.2395 |
| 2084,H | 0.2388 | 0.0100 | 0.1987 | 0.0174 | 0.1864 | 0.0153 | 0.1878 | 0.0093 |
| 2085,C | -0.4879 | 0.0884 | -0.5794 | 0.2947 | -0.3391 | 0.2985 | -0.0530 | 0.1858 |
| 2086,N | -0.1489 | 0.0238 | -0.2020 | 0.1088 | -0.1693 | 0.0247 | -0.0763 | 0.0480 |
| 2087,O | -0.1314 | 0.0416 | -0.1411 | 0.0309 | -0.1745 | 0.0362 | -0.1454 | 0.0647 |
| 2088,O | -0.0902 | 0.0261 | -0.1244 | 0.0407 | -0.1667 | 0.0278 | -0.1579 | 0.0624 |

**Supplementary Table 10.** Harmonic vibrational frequency calculations for all optimized structures.

| Frame # | Raw reactant total energy | Raw TS total energy | Raw product total energy | Relative reactant energy (kcal/mol) | Relative TS energy (kcal/mol) | Relative product energy (kcal/mol) |
|---|---|---|---|---|---|---|
| 1 | −1930.584 | −1930.551 | −1930.623 | 0.000 | 21.039 | −24.052 |
| 2 | −1922.338 | −1922.317 | −1922.366 | 0.000 | 13.400 | −17.865 |
| 3 | −1931.135 | −1931.135 | −1931.171 | 0.000 | 0.103 | −22.734 |
| 4 | −1926.903 | −1926.875 | −1926.941 | 0.000 | 17.703 | −23.645 |
| 5 | −1935.147 | −1935.114 | −1935.173 | 0.000 | 20.756 | −16.304 |
| **Averaged** | | | | **0.000** | **14.600** | **−20.920** |

**Supplementary Table 11.** Gas Phase QM region data.

| Frame # | Raw reactant total energy | Raw TS total energy | Raw product total energy | Relative reactant energy (kcal/mol) | Relative TS energy (kcal/mol) | Relative product energy (kcal/mol) |
|---|---|---|---|---|---|---|
| 1 | −1932.998 | −1932.958 | −1933.029 | 0.000 | 25.095 | −19.971 |
| 2 | −1932.982 | −1932.947 | −1933.017 | 0.000 | 22.133 | −21.981 |
| 3 | −1932.994 | −1932.952 | −1933.026 | 0.000 | 26.309 | −20.553 |
| 4 | −1933.003 | −1932.970 | −1933.037 | 0.000 | 20.496 | −21.868 |
| 5 | −1932.997 | −1932.965 | −1933.030 | 0.000 | 19.803 | −20.700 |
| **Averaged** | | | | **0.000** | **22.767** | **−21.015** |

**Supplementary Table 12.** Comparison of experimental and calculated NMR chemical shifts for Tyr9.

| Atom (Tyr9) | State | Experimental (ppm) | Calculated (ppm) | Δ(Calc-Exp) (ppm) |
|---|---|---|---|---|
| HE2 | Free | 6.574 | 6.787 | 0.033 |
| CE2 | Free | 118.585 | 118.682 | 0.097 |
| HE2 | TSA–bound | 7.242 | 7.160 | −0.082 |
| CE2 | TSA–bound | 119.670 | 119.905 | 0.235 |

**Supplementary Table 13.** Structural statistics over 10 lowest-energy NMR structures of free (PDB: 9SMT) and 6-NBT–bound KABLE2.5 (PDB: 29SB). Average root-mean-square deviation from the mean structure calculated for residues 2-125.

| | Free KABLE2.5 (PDB: 9SMT) | 6-NBT–bound KABLE2.5 (PDB: 29SB) |
|---|---|---|
| **NMR distance and dihedral constraints** | | |
| Distance constraints | 3600 | 3620 |
| Total NOE | | |
| Intra-residue | | |
| Inter-residue | | |
| Sequential (\|i – j\| = 1) | 1423 | 1423 |
| Medium-range (\|i – j\| < 4) | 1117 | 1117 |
| Long-range (\|i – j\| > 5) | 1060 | 1060 |
| Intermolecular | | 20 |
| Hydrogen bonds | | |
| Total dihedral angle restraints | 237 | 237 |
| ϕ | | |
| ψ | | |
| **Structure statistics** | | |
| Violations (mean and s.d.) | | |
| Distance constraints (Å) | | |
| Dihedral angle constraints (°) | | |
| Max. dihedral angle violation (°) | 0 | 0.04 |
| Max. distance constraint violation (Å) | 0 | 0 |
| Deviations from idealized geometry | | |
| Bond lengths (Å) | | |
| Bond angles (°) | | |
| Impropers (°) | | |
| Average pairwise r.m.s. deviation** (Å) | | |
| Heavy | 0.77 ± 0.04 | 1.20 ± 0.06 |
| Backbone | 0.22 ± 0.03 | 0.52 ± 0.09 |

**Pairwise r.m.s. deviation was calculated among 10 refined structures

**Supplementary References.**


1 Gadekar, S. C. *et al.* Rerouting the Organocatalytic Benzoin Reaction toward Aldehyde Deuteration. *ACS Catal.* **11**, 14561-14569 (2021).

2 Kemp, D. S. & Woodward, R. B. The N-ethylbenzisoxazolium cation—I: Preparation and reactions with nucleophilic species. *Tetrahedron* **21**, 3019-3035 (1965).

3 Hollfelder, F., Kirby, A. J., Tawfik, D. S., Kikuchi, K. & Hilvert, D. Characterization of Proton-Transfer Catalysis by Serum Albumins. *J. Am. Chem. Soc.* **122**, 1022-1029 (2000).

4 Manetsch, R., Zheng, L., Reymond, M. T., Woggon, W. D. & Reymond, J. L. A catalytic antibody against a tocopherol cyclase inhibitor. *Chem. Eur. J.* **10**, 2487-2506 (2004).

5 Frisch, M. J. *et al.* Gaussian 16 Revision B.01. (2016).

6 Delaglio, F. *et al.* NMRPipe: A multidimensional spectral processing system based on UNIX pipes. *J. Biomol. NMR* **6**, 277-293 (1995).

7 Vranken, W. F. *et al.* The CCPN data model for NMR spectroscopy: Development of a software pipeline. *Proteins* **59**, 687-696 (2005).

8 Chen, Y. *et al.* Emergence of specific binding and catalysis from a designed generalist binding protein. *Nat. Chem.* **18**, 1334-1344 (2026).

9 Larina, L. I. & Milata, V. 1H, 13C and 15N NMR spectroscopy and tautomerism of nitrobenzotriazoles. *Magn. Reson. Chem.* **47**, 142-148 (2009).

10 Cheung, M. S., Maguire, M. L., Stevens, T. J. & Broadhurst, R. W. DANGLE: A Bayesian inferential method for predicting protein backbone dihedral angles and secondary structure. *J. Magn. Reson.* **202**, 223-233 (2010).

11 Güntert, P. & Buchner, L. Combined automated NOE assignment and structure calculation with CYANA. *J. Biomol. NMR* **62**, 453-471 (2015).

12 Brünger, A. T. *et al.* Crystallography & NMR system: A new software suite for macromolecular structure determination. *Acta Crystallogr. D Biol. Crystallogr.* **54**, 905-921 (1998).

13 Schwieters, C. D., Kuszewski, J. J., Tjandra, N. & Clore, G. M. The Xplor-NIH NMR molecular structure determination package. *J. Magn. Reson.* **160**, 65-73 (2003).

14 Caselle, E. A. *et al.* Kemp Eliminases of the AlleyCat Family Possess High Substrate Promiscuity. *ChemCatChem* **11**, 1425-1430 (2019).

15 Hwang, T. L., van Zijl, P. C. & Mori, S. Accurate quantitation of water-amide proton exchange rates using the phase-modulated CLEAN chemical EXchange (CLEANEX-PM) approach with a Fast-HSQC (FHSQC) detection scheme. *J. Biomol. NMR* **11**, 221-226 (1998).

16 Skinner, J. J., Lim, W. K., Bédard, S., Black, B. E. & Englander, S. W. Protein hydrogen exchange: Testing current models. *Protein Sci.* **21**, 987-995 (2012).
